# Transcriptomic profiling reveals estradiol and progesterone specific microRNA signatures differentially regulate mRNA targets in vaginal epithelial cells

**DOI:** 10.64898/2026.09.17.752383

**Authors:** Shreya Joshi, Aisha Nazli, Chris Verschoor, Charu Kaushic

**Affiliations:** McMaster Immunology Research Centre, Michael G. DeGroote Centre for Learning and Discovery, McMaster University, Hamilton, ON, Canada; Department of Medicine, McMaster University, Hamilton, ON, Canada; Health Sciences North Research Institute, Sudbury, ON, Canada; NOSM University, Sudbury, ON, Canada

**Keywords:** Vaginal epithelial cells, estradiol (E2), progesterone (P4), microRNAs (miRNAs/miRs), epithelial barrier, sex hormones

## Abstract

Epithelial cells of the female reproductive tract (FRT) form a protective barrier and serve as the first line of defense against sexually transmitted infections (STIs). They recognize pathogens, release cytokines and initiate innate immune responses. Sex hormones 17β-estradiol (E2) and progesterone (P4) fluctuate throughout the menstrual cycle and influence epithelial cell functions. Generally, E2 confers protection against viral STIs like HIV-1 and HSV-2, whereas P4 increases susceptibility. However, the underlying mechanisms remain incompletely understood. MicroRNAs, small non-coding RNAs that regulate gene expression by inhibiting translation of target mRNAs, significantly influence viral infections, by promoting or restricting viral pathogenesis. Here, we examined the effect of sex hormones on microRNA expression in vaginal epithelial cells (VECs), the first responders to pathogens in the FRT. Using next generation sequencing analysis, the effect of physiological concentrations of E2 or P4 on microRNA transcription profile of VECs grown in air-liquid interface cultures was characterized. Thirteen miRNAs were uniquely regulated by E2 and 102 by P4, with 29 miRNAs that overlapped between both. The top four differentially expressed miRNAs from both treatments were validated by RT-qPCR. The mRNA targets of unique E2-miRNAs were involved in adherens junction, tight junction and chemokine signaling pathways; whereas the mRNA targets of unique P4-miRNAs were involved in MAPK signaling pathway. Interestingly, several miRNAs regulated by E2-(eg. hsa-miR-9-3p, hsa-miR-200c-5p, hsa-miR-548c-5p) and by P4 (eg. hsa-let-7f-5p, hsa-miR-16-5p, hsa-miR-27b-3p) targeted genes related to cell cycle pathway. Together, these findings indicate that microRNAs regulated by E2 and P4 influence mRNAs in the pathways associated with cell adhesion, cell growth and development and innate immune responses, critical for maintenance of epithelial barrier integrity and regulating pathogen susceptibility in the vaginal tract. These hormone-regulated miRNAs may serve as potential targets for therapeutic interventions.

## 1. Introduction

The mucosal epithelial cells of the female reproductive tract (FRT) act as a protective physical barrier and constitute the first line of defense against sexually transmitted pathogens, including bacteria, viruses and fungi. They also act as immune sentinels by recognizing pathogen specific structures, producing antimicrobial proteins and releasing cytokines and chemokines that recruit immune cells at the site of infection and influence adaptive responses^[1]^.

The FRT is divided into two distinct anatomical sites- the upper and the lower FRT. The upper FRT consisting of the fallopian tubes, endometrium and endocervix, is lined by single layer of columnar epithelial cells. In contrast, the lower FRT consisting of the ectocervix and vagina, is lined by stratified squamous epithelial cells that undergo constant proliferation, forming a multilayered protective barrier against invading microbes^[1–3]^. The continuous shedding of the apical layer of cells also prevents microbial colonization and barrier breach^[4–6]^. Various intercellular junctions mainly present in the basal layers of the lower FRT also play a crucial role in maintaining epithelial organization, barrier integrity and prevention of pathogen entry^[5]^. Tight junctions (ZO-1, JAM-A and claudin-1), adherens junctions (E-cadherin) and desmosomes (JAM3) are the most abundant cell-cell adhesions present in the lower FRT^[7]^. The epithelial cells express a wide range of pattern recognition receptors such as toll like receptors (TLRs) that detect an array of pathogen-associated molecular patterns and elicit an innate immune response^[6, 8]^. It activates intracellular signaling cascades resulting in the production of pro-inflammatory cytokines such as interleukin 6 (IL-6), tumor necrosis factor α (TNF-α) and interferons; chemokines such as IL-8, macrophage inflammatory protein 1α (MIP-1α) and MIP-1β that recruit other immune cells and promote pathogen clearance^[9]^. In addition, they also produce antimicrobial peptides such as defensins, lysozyme, secretory leucocyte protease inhibitors and lactoferrin, both constitutively and in response to infection, providing protection against invading pathogens^[1, 10]^.

Sex hormones, 17β-estradiol (E2) and progesterone (P4) regulate the physiology and functions of the FRT epithelial cells. Fluctuations in these hormones during the menstrual cycle affect epithelial barrier integrity and innate immune functions. E2 enhances the expression of cell junction proteins and reduces inflammation, whereas P4 exerts opposite effect by reducing the expression of junction proteins and increasing inflammation^[8, 11]^. These hormone-mediated effects significantly influence susceptibility to sexually transmitted infections (STIs), including Chlamydia, Gonorrhoea, HIV-1 and HSV-2 in women^[3, 12]^. For example, work done in *ex-vivo* tissue culture models, clinical studies, and non-human primates suggests that women in E2-high follicular phase of the menstrual cycle are better protected against HIV-1 infection whereas more prone to acquiring HIV-1 during the P4-high luteal phase^[13–16]^. However, the molecular mechanisms underlying the hormone mediated differential protection or susceptibility to STIs are not known.

MicroRNAs (miRNAs/miRs) are small, non-coding, single stranded RNAs playing a significant role in post transcriptional regulation of gene expression. Typically, they interact with 3’UTR of target mRNAs leading to translation inhibition or mRNA cleavage ^[17–19]^. MiRNAs regulate critical biological processes such as cell proliferation, differentiation, apoptosis, metabolism, and their aberrant expression can influence disease outcomes^[20, 21]^. Studies have shown that sex hormones regulate microRNA expression in various hormone-responsive tissues including the endometrium in the FRT^[22–25]^. However, there is no report yet of hormone-mediated microRNA regulation in vaginal epithelial cells. Given the large surface area of the vagina and it being the first mucosal surface to come into contact with sexually transmitted pathogens, characterizing the sex hormone-mediated global microRNA profile of vaginal epithelial cells is important^[4, 26]^.

Here, for the first time, we used next generation sequencing (NGS) to characterize the global microRNA transcriptional response of vaginal epithelial cells grown in air-liquid-interface (ALI) cultures in the presence or absence of physiological levels of female sex hormones, E2 and P4. We report that 13 miRNAs were regulated by E2 and 102 miRNAs by P4, along with 29 miRNAs that overlapped between both treatments. The mRNA targets of unique miRNAs identified in E2- and P4-treated VK2 cells were involved in pathways related to epithelial barrier integrity, cell growth and development, and innate immune responses, crucial for protection against STIs.

## 2. Materials and Methods

### 2.1 VK2 cell preparation and ALI cultures

VK2/E6E7 vaginal epithelial cells (ATCC-CRL 2616) were used to generate ALI cultures, as described previously^[27]^. Briefly, 60,000 VK2 cells were seeded in transwell polystyrene inserts with a pore size of 0.4 μm (BD Falcon, Mississauga, ON, Canada). Keratinocyte serum free media (KSFM) (Cat.# 17005042, Thermofisher) supplemented with 0.1 ng/ml of human recombinant epidermal growth factor (EGF), 0.05 mg/ml bovine pituitary extract (BPE), 0.4 Mm CaCl2 and 100 units/ml penicillin/streptomycin (Sigma Aldrich, Oakville, ON, Canada) was added to the apical and basolateral side of the cultures. To create ALI cultures, media from the apical side was removed 24 hours after seeding cells.

### 2.2 Hormone treatment of VK2 cell cultures

VK2 cells were grown for 6 days in the presence or absence of E2 (10^-9^M; Sigma-Aldrich), P4 (10^-7^M; Sigma-Aldrich) or no hormone (NH) as described earlier^[5]^. The concentrations of E2 and P4 correspond to the highest serum levels during menstrual cycle^[28]^. Both hormones were prepared in KSFM and the basolateral hormone containing media was replenished every 48 hours for 6 days.

### 2.3 MicroRNA extraction and screening by Next Generation Sequencing (NGS)

Total RNA including microRNAs was extracted using miRNeasy Tissue/Cells Advanced Mini Kit (Cat# 217604, Qiagen, Canada) according to manufacturer’s protocol. The purified RNA was resuspended in RNase-free water and was quantified using the Nanodrop One spectrophotometer (Thermo Scientific, Waltham, MA). RNA samples were extracted and analyzed separately from four different experiments (n=4) performed on different days.

All samples were sequenced at the London Regional Genomics Centre (Robarts Research Institute, London, Ontario, Canada; http://www.lrgc.ca) using the Illumina NextSeq 500 (Illumina Inc., San Diego, CA). Total RNA samples were quantified using the NanoDrop (Thermo Fisher Scientific, Waltham, MA) and quality was assessed using 1 uL (50-500 ng/uL) of sample on the Agilent 4150 TapeStation and RNA Screen Tape (Agilent Technologies Inc., Palo Alto, CA). They were then processed using the QIAseq miRNA Library Kit (Qiagen, Carlsbad, CA). Briefly, samples underwent ligation, cDNA synthesis, amplification with indexed primers, and clean-up. Libraries were then quantitated using the Qubit 2.0 Fluorimeter (Thermo Fisher Scientific, Waltham, MA) and equimolar pooled into one library. Size distribution was assessed on an Agilent 4150 TapeStation High Sensitivity DNA Screen Tape. The library was sequenced on an Illumina NextSeq 500 as 76 bp single end run, using one High Output v2 kit (75 cycles).

### 2.4 Identification of Differentially Expressed (DE)-miRNAs

Pre-processing of the raw data was performed on the Galaxy platform online and miRNA raw counts were obtained^[29]^. They were loaded into R v4.4.1 and experimental batch effects (n=4) were removed using the function Combat_seq in ‘sva’ package^[30]^. Following batch correction, differential expression analysis for E2 vs NH and P4 vs NH comparisons was performed using the package ‘DESeq2’ in R^[31]^. The Benjamini-Hochberg procedure with a false discovery rate (FDR) threshold of <0.05 was used to adjust p-values for multiple testing. However, this approach yielded no significant miRNAs for downstream analysis in E2 and 65 in P4. Even when applying an adjusted p < 0.15 threshold, we did not identify any significant miRNAs in E2, and 117 miRNAs were detected in P4. Hence, a less stringent, unadjusted p < 0.05 significance threshold was used for downstream analysis, for both E2 and P4 treatments. To ensure that this did not lead to false or non-relevant miRNAs, validation was done by RT-qPCR.

### 2.5 Validation of sequencing data by quantitative real-time PCR

Quantitative real-time PCR (RT-qPCR) was performed to validate miRNAs regulated by E2 and P4 identified in sequencing analysis. RNA was extracted as described in methods 2.3. cDNA was synthesized using Qiagen’s miRCURY LNA RT kit (Cat.# 339340) and RT-qPCR was performed using miRCURY LNA SYBR Green PCR Kit (Qiagen, Cat.# 339345) on Quant Studio 3 real-time PCR system (Thermo Fisher Scientific). miRCURY LNA miRNA PCR Assays (Qiagen, Cat.#. 339306) were used for validation and their Gene Globe IDs are listed in supplementary table 1. Samples were run in duplicate and all data was normalized to hsa-miR-103a-3p as an endogenous control identified using RefFinder^[32]^. GraphPad Prism v8.0.1 (GraphPad Software, San Diego, CA, USA) was used for statistical analysis.

### 2.6 mRNA Target Prediction and Pathway Enrichment Analysis

The unique differentially expressed miRNAs in E2- and P4-treated VK2 cells (unadjusted p<0.05) were loaded onto miRNet 2.0, a web-based platform to investigate miRNA-mRNA regulatory networks^[33]^. Only experimentally validated, high confidence mRNA targets were predicted using miRTarBase v9.0^[34]^. The resulting interaction tables were used to create and visualize miRNA-mRNA networks. Because of the large P4-miRNA-mRNA network containing 9240 target mRNAs, a degree filter of 10 was applied to condense the network and reduce its complexity. Following this, the mRNA targets were subjected to pathway enrichment analysis using the KEGG (Kyoto Encyclopedia of Genes and Genomes) database in miRNet to identify biological pathways regulated by the target mRNAs. Enrichment was performed using a hypergeometric over-representation analysis and adjusted for multiple testing using Benjamini-Hochberg method. Signaling pathways with FDR <0.05 were considered statistically significant. Given the propensity for ubiquitous signaling pathways to be enriched by chance, we performed a KEGG pathway enrichment analysis on the mRNA targets of 20 random miRNAs from the differential expression dataset (excluding E2 and P4 regulated miRNAs) and 51 pathways were identified (Table 1). Pathways found to be significant (FDR < 0.05; highlighted in yellow in Table 1) were subsequently removed from E2 and P4 DE-miRNA pathway analysis, to ensure that there were no false or non-relevant pathways.

**Table 1:**
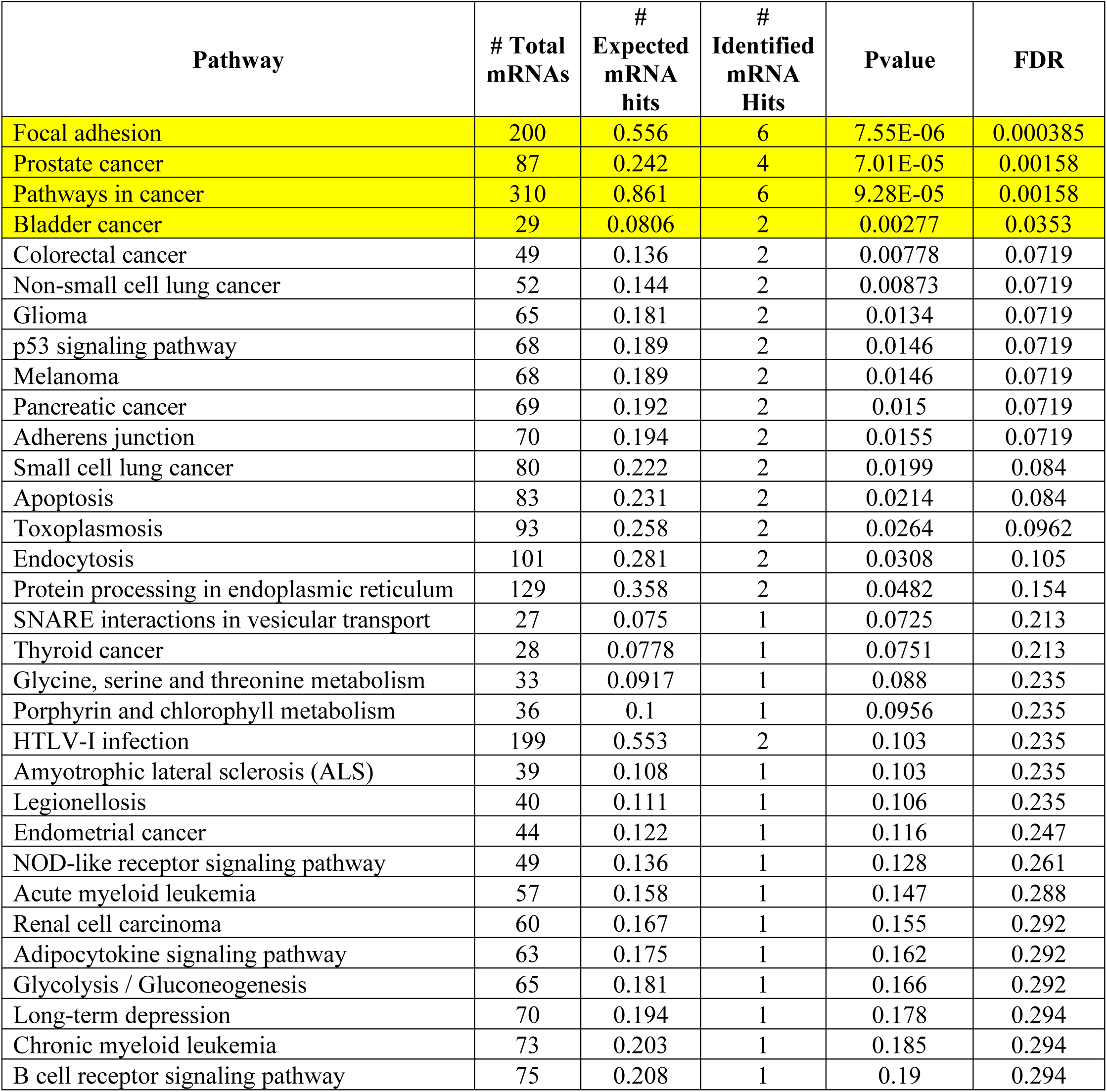
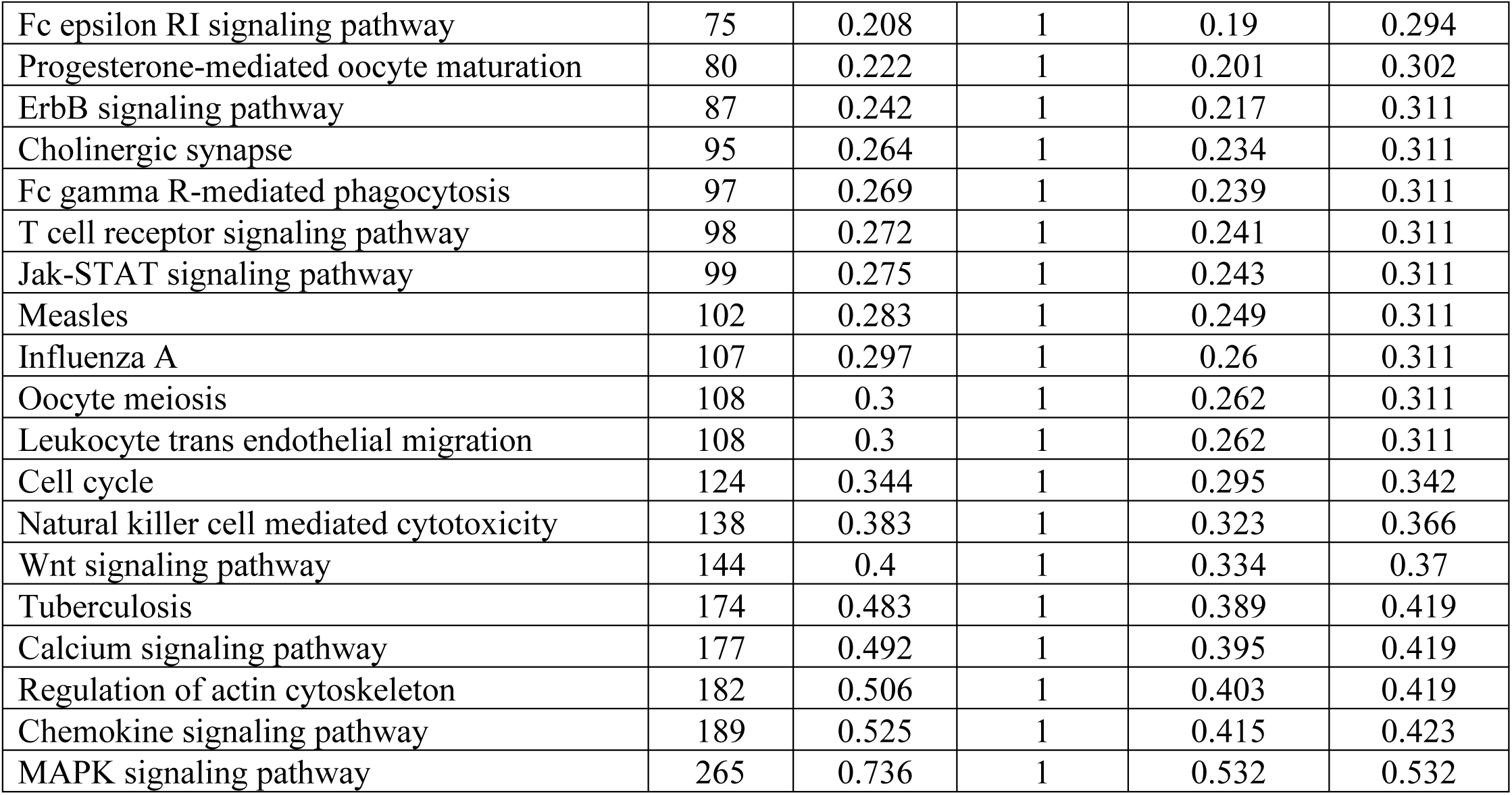
KEGG pathway enrichment of mRNA targets of 20 random miRNAs from differential expression dataset (excluding E2- and P4-regulated miRNAs). Pathways highlighted in yellow are statistically significant (FDR < 0.05)

| Pathway | # Total mRNAs | # Expected mRNA hits | # Identified mRNA Hits | Pvalue | FDR |
| --- | --- | --- | --- | --- | --- |
| Focal adhesion | 200 | 0.556 | 6 | 7.55E-06 | 0.000385 |
| Prostate cancer | 87 | 0.242 | 4 | 7.01E-05 | 0.00158 |
| Pathways in cancer | 310 | 0.861 | 6 | 9.28E-05 | 0.00158 |
| Bladder cancer | 29 | 0.0806 | 2 | 0.00277 | 0.0353 |
| Colorectal cancer | 49 | 0.136 | 2 | 0.00778 | 0.0719 |
| Non-small cell lung cancer | 52 | 0.144 | 2 | 0.00873 | 0.0719 |
| Glioma | 65 | 0.181 | 2 | 0.0134 | 0.0719 |
| p53 signaling pathway | 68 | 0.189 | 2 | 0.0146 | 0.0719 |
| Melanoma | 68 | 0.189 | 2 | 0.0146 | 0.0719 |
| Pancreatic cancer | 69 | 0.192 | 2 | 0.015 | 0.0719 |
| Adherens junction | 70 | 0.194 | 2 | 0.0155 | 0.0719 |
| Small cell lung cancer | 80 | 0.222 | 2 | 0.0199 | 0.084 |
| Apoptosis | 83 | 0.231 | 2 | 0.0214 | 0.084 |
| Toxoplasmosis | 93 | 0.258 | 2 | 0.0264 | 0.0962 |
| Endocytosis | 101 | 0.281 | 2 | 0.0308 | 0.105 |
| Protein processing in endoplasmic reticulum | 129 | 0.358 | 2 | 0.0482 | 0.154 |
| SNARE interactions in vesicular transport | 27 | 0.075 | 1 | 0.0725 | 0.213 |
| Thyroid cancer | 28 | 0.0778 | 1 | 0.0751 | 0.213 |
| Glycine, serine and threonine metabolism | 33 | 0.0917 | 1 | 0.088 | 0.235 |
| Porphyrin and chlorophyll metabolism | 36 | 0.1 | 1 | 0.0956 | 0.235 |
| HTLV-I infection | 199 | 0.553 | 2 | 0.103 | 0.235 |
| Amyotrophic lateral sclerosis (ALS) | 39 | 0.108 | 1 | 0.103 | 0.235 |
| Legionellosis | 40 | 0.111 | 1 | 0.106 | 0.235 |
| Endometrial cancer | 44 | 0.122 | 1 | 0.116 | 0.247 |
| NOD-like receptor signaling pathway | 49 | 0.136 | 1 | 0.128 | 0.261 |
| Acute myeloid leukemia | 57 | 0.158 | 1 | 0.147 | 0.288 |
| Renal cell carcinoma | 60 | 0.167 | 1 | 0.155 | 0.292 |
| Adipocytokine signaling pathway | 63 | 0.175 | 1 | 0.162 | 0.292 |
| Glycolysis / Gluconeogenesis | 65 | 0.181 | 1 | 0.166 | 0.292 |
| Long-term depression | 70 | 0.194 | 1 | 0.178 | 0.294 |
| Chronic myeloid leukemia | 73 | 0.203 | 1 | 0.185 | 0.294 |
| B cell receptor signaling pathway | 75 | 0.208 | 1 | 0.19 | 0.294 |
| Fc epsilon RI signaling pathway | 75 | 0.208 | 1 | 0.19 | 0.294 |
| Progesterone-mediated oocyte maturation | 80 | 0.222 | 1 | 0.201 | 0.302 |
| ErbB signaling pathway | 87 | 0.242 | 1 | 0.217 | 0.311 |
| Cholinergic synapse | 95 | 0.264 | 1 | 0.234 | 0.311 |
| Fc gamma R-mediated phagocytosis | 97 | 0.269 | 1 | 0.239 | 0.311 |
| T cell receptor signaling pathway | 98 | 0.272 | 1 | 0.241 | 0.311 |
| Jak-STAT signaling pathway | 99 | 0.275 | 1 | 0.243 | 0.311 |
| Measles | 102 | 0.283 | 1 | 0.249 | 0.311 |
| Influenza A | 107 | 0.297 | 1 | 0.26 | 0.311 |
| Oocyte meiosis | 108 | 0.3 | 1 | 0.262 | 0.311 |
| Leukocyte trans endothelial migration | 108 | 0.3 | 1 | 0.262 | 0.311 |
| Cell cycle | 124 | 0.344 | 1 | 0.295 | 0.342 |
| Natural killer cell mediated cytotoxicity | 138 | 0.383 | 1 | 0.323 | 0.366 |
| Wnt signaling pathway | 144 | 0.4 | 1 | 0.334 | 0.37 |
| Tuberculosis | 174 | 0.483 | 1 | 0.389 | 0.419 |
| Calcium signaling pathway | 177 | 0.492 | 1 | 0.395 | 0.419 |
| Regulation of actin cytoskeleton | 182 | 0.506 | 1 | 0.403 | 0.419 |
| Chemokine signaling pathway | 189 | 0.525 | 1 | 0.415 | 0.423 |
| MAPK signaling pathway | 265 | 0.736 | 1 | 0.532 | 0.532 |

## 3. Results

### 3.1 E2 and P4 significantly regulate microRNA expression in VK2 cells

To determine the effect of E2 and P4 on the microRNA transcriptional profile in vaginal epithelial cells, VK2 cells were grown in the presence or absence of physiological levels of E2 (10^−9^ M; n=4), P4 (10^−7^ M; n=4) or NH (n=4), in ALI cultures. At the end of culture on day 6, RNA was extracted from the cells and microRNA NGS was performed. A multidimensional scaling (MDS) was conducted on the top 100 most variable miRNAs to assess overall similarities in the transcriptional profiles of E2, P4 and NH-treated VK2 cells. Samples from each treatment group formed separate clusters in the MDS plot (Figure 1A). To evaluate whether the miRNA expression profiles were significantly different with each treatment compared to NH, a pairwise permutational multivariate analysis of variance (PERMANOVA) was performed in R. While E2 was not significant (p = 0.28), it still contributes 15% of the total variance in miRNA expression (R^2^ = 0.15). P4 can be concluded to significantly change the overall miRNA expression profile (p = 0.03) and accounts for 25% of the total variance (R^2^ = 0.25). Differentially expressed microRNAs (DE-miRNAs) meeting the conservative adjusted p < 0.05 and adjusted p < 0.15 thresholds were not identified in VK2 cells treated with E2, whereas 65 and 117 miRNAs were identified in P4-treated VK2 cells under these respective thresholds. Thus, a less stringent, unadjusted p < 0.05, transcriptomic profiling was used for both E2 and P4. E2-treated VK2 cells had 42 DE-miRNAs (supplementary table 2). Of these, 24 miRNAs were upregulated with 10 showing strong upregulation (log2FC > 0.5) (Table 2, Figure 1B), whereas 18 miRNAs were downregulated, 4 of which showed strong downregulation (log2FC < -0.5) (Table 2, Figure 1B). In P4 treated VK2 cells, 131 DE-miRNAs were identified (supplementary table 3). Among them, 76 miRNAs were upregulated with 25 showing strong upregulation (log2FC > 0.5) (Table 3, Figure 1C); and 55 miRNAs were downregulated with 20 showing strong downregulation (log2FC < -0.5) (Table 3, Figure 1C).

**Figure 1:**
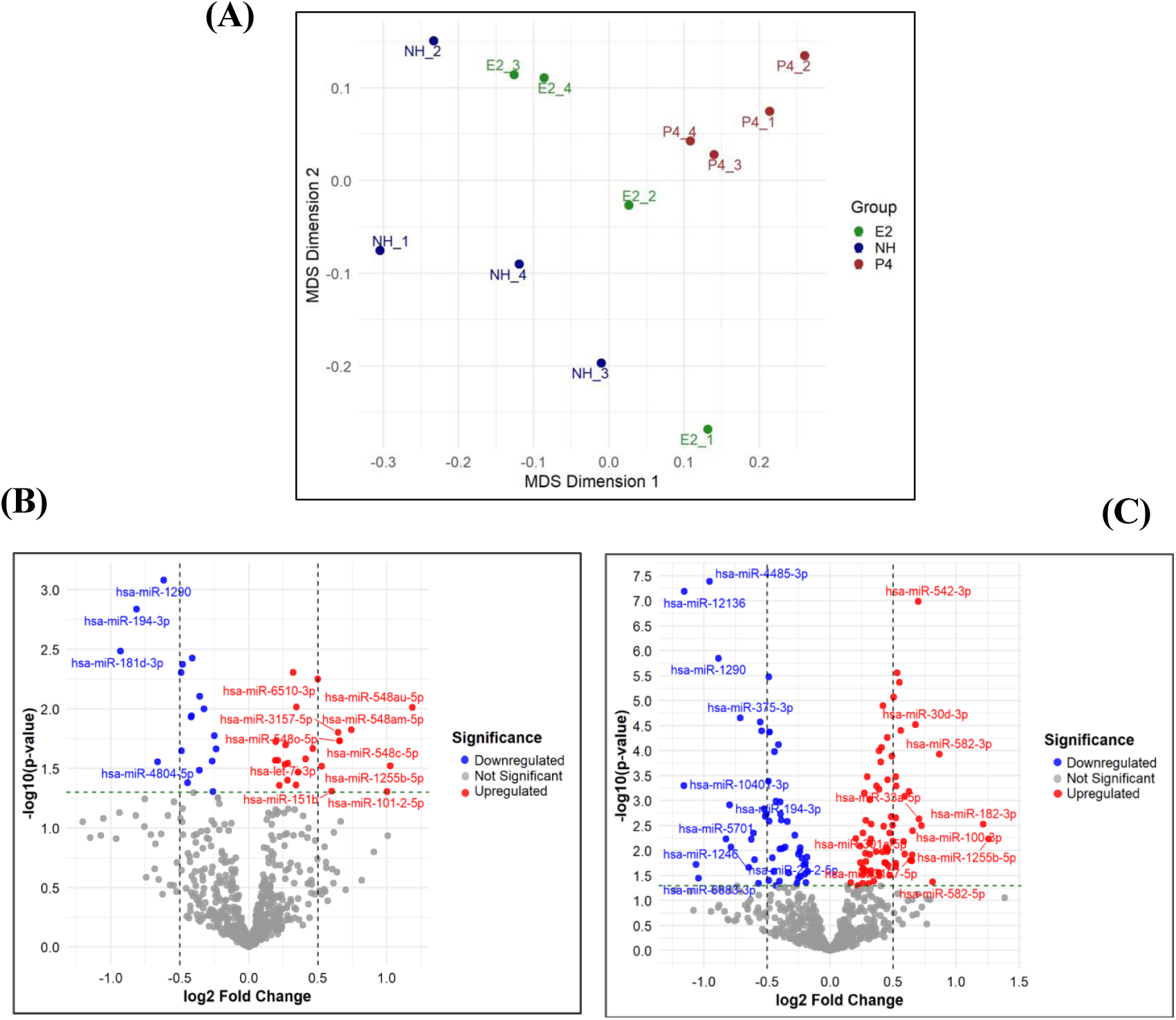
E2 and P4 significantly regulate microRNA expression in VK2 cells. (A) MDS plot of top 100 most variable miRNAs in E2, P4 and NH treated VK2 cells (n=4, each). (B)Volcano plot showing E2-DE-miRNAs relative to NH in VK2 cells. The horizontal green dashed line marks the significance threshold (unadjusted p < 0.05). Red and blue dots represent significantly up- and down-regulated miRNAs, respectively; and gray dots represent non-significant miRNAs. The miRNAs with strong differential expression (|log2FC>0.5|), indicated by the vertical black line, are labelled. (C) Volcano plot showing P4-DE-miRNAs relative to NH in VK2 cells. The horizontal green dashed line marks the significance threshold (unadjusted p < 0.05). Red and blue dots represent significantly up- and down-regulated miRNAs, respectively; and gray dots represent non-significant miRNAs. The top 10 strongly up and downregulated miRNAs (|log2FC>0.5|), marked by the vertical black line are labelled.

**Table 2:**
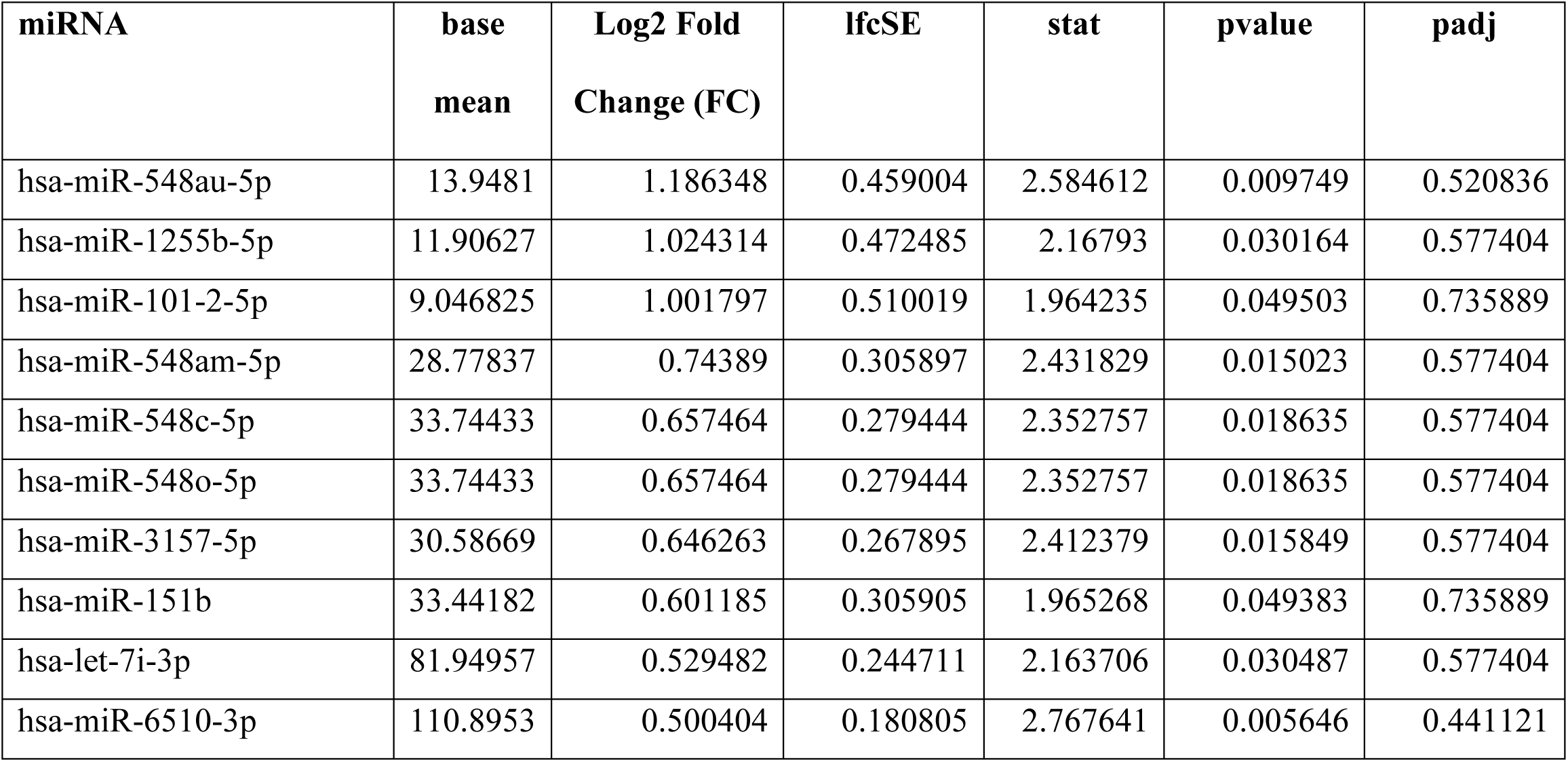
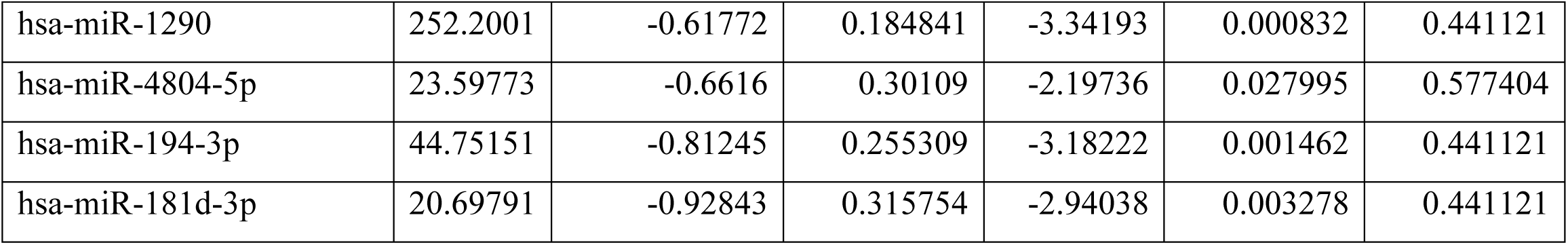
List of E2-DE-miRNAs showing strong upregulation (unadjusted p < 0.05, log2FC > 0.5) and downregulation (unadjusted p < 0.05, log2FC < -0.5) in VK2 cells.

| miRNA | base<br>mean | Log2 Fold<br>Change (FC) | lfcSE | stat | pvalue | padj |
| --- | --- | --- | --- | --- | --- | --- |
| hsa-miR-548au-5p | 13.9481 | 1.186348 | 0.459004 | 2.584612 | 0.009749 | 0.520836 |
| hsa-miR-1255b-5p | 11.90627 | 1.024314 | 0.472485 | 2.16793 | 0.030164 | 0.577404 |
| hsa-miR-101-2-5p | 9.046825 | 1.001797 | 0.510019 | 1.964235 | 0.049503 | 0.735889 |
| hsa-miR-548am-5p | 28.77837 | 0.74389 | 0.305897 | 2.431829 | 0.015023 | 0.577404 |
| hsa-miR-548c-5p | 33.74433 | 0.657464 | 0.279444 | 2.352757 | 0.018635 | 0.577404 |
| hsa-miR-548o-5p | 33.74433 | 0.657464 | 0.279444 | 2.352757 | 0.018635 | 0.577404 |
| hsa-miR-3157-5p | 30.58669 | 0.646263 | 0.267895 | 2.412379 | 0.015849 | 0.577404 |
| hsa-miR-151b | 33.44182 | 0.601185 | 0.305905 | 1.965268 | 0.049383 | 0.735889 |
| hsa-let-7i-3p | 81.94957 | 0.529482 | 0.244711 | 2.163706 | 0.030487 | 0.577404 |
| hsa-miR-6510-3p | 110.8953 | 0.500404 | 0.180805 | 2.767641 | 0.005646 | 0.441121 |
| hsa-miR-1290 | 252.2001 | -0.61772 | 0.184841 | -3.34193 | 0.000832 | 0.441121 |
| hsa-miR-4804-5p | 23.59773 | -0.6616 | 0.30109 | -2.19736 | 0.027995 | 0.577404 |
| hsa-miR-194-3p | 44.75151 | -0.81245 | 0.255309 | -3.18222 | 0.001462 | 0.441121 |
| hsa-miR-181d-3p | 20.69791 | -0.92843 | 0.315754 | -2.94038 | 0.003278 | 0.441121 |

**Table 3:**
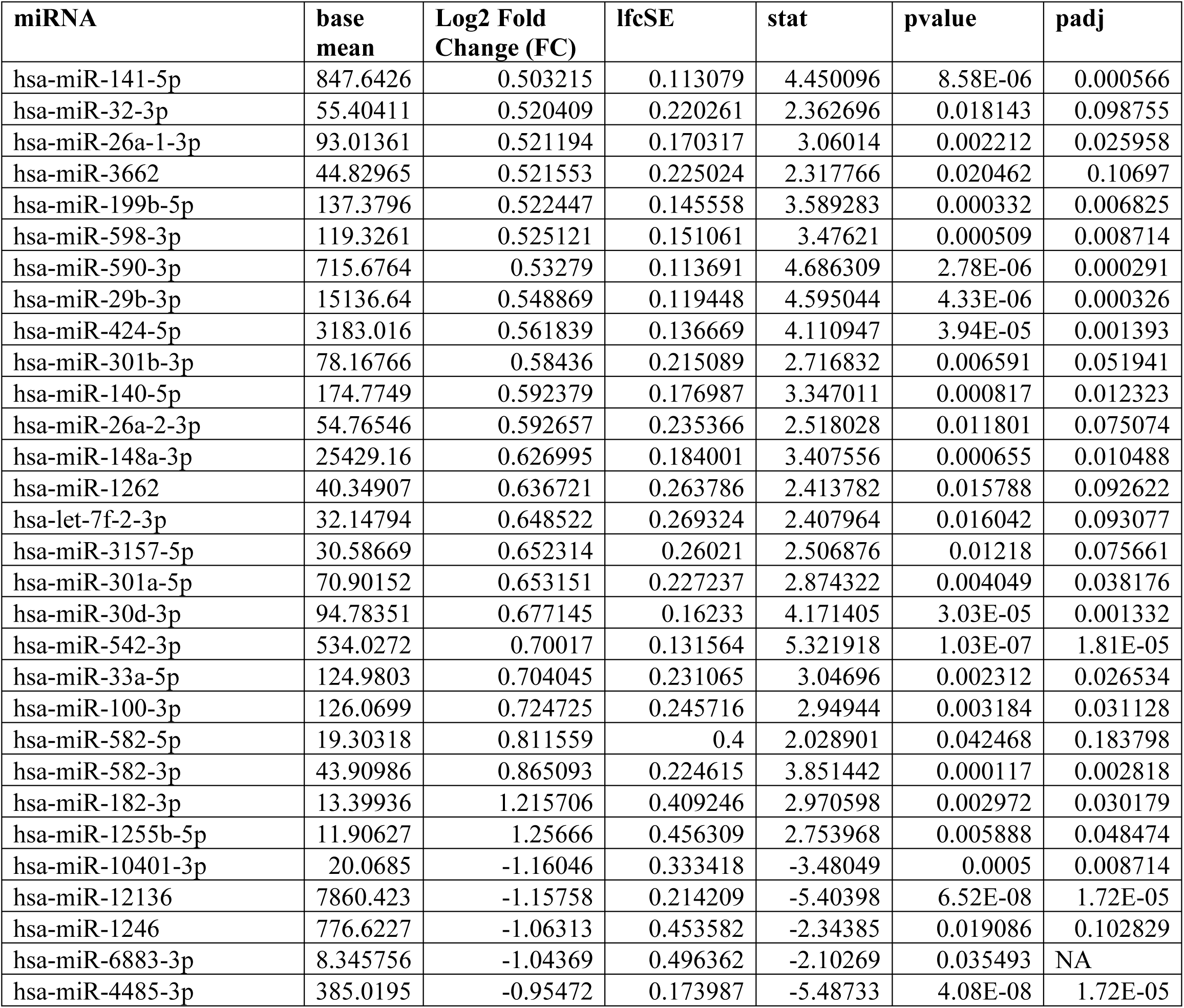
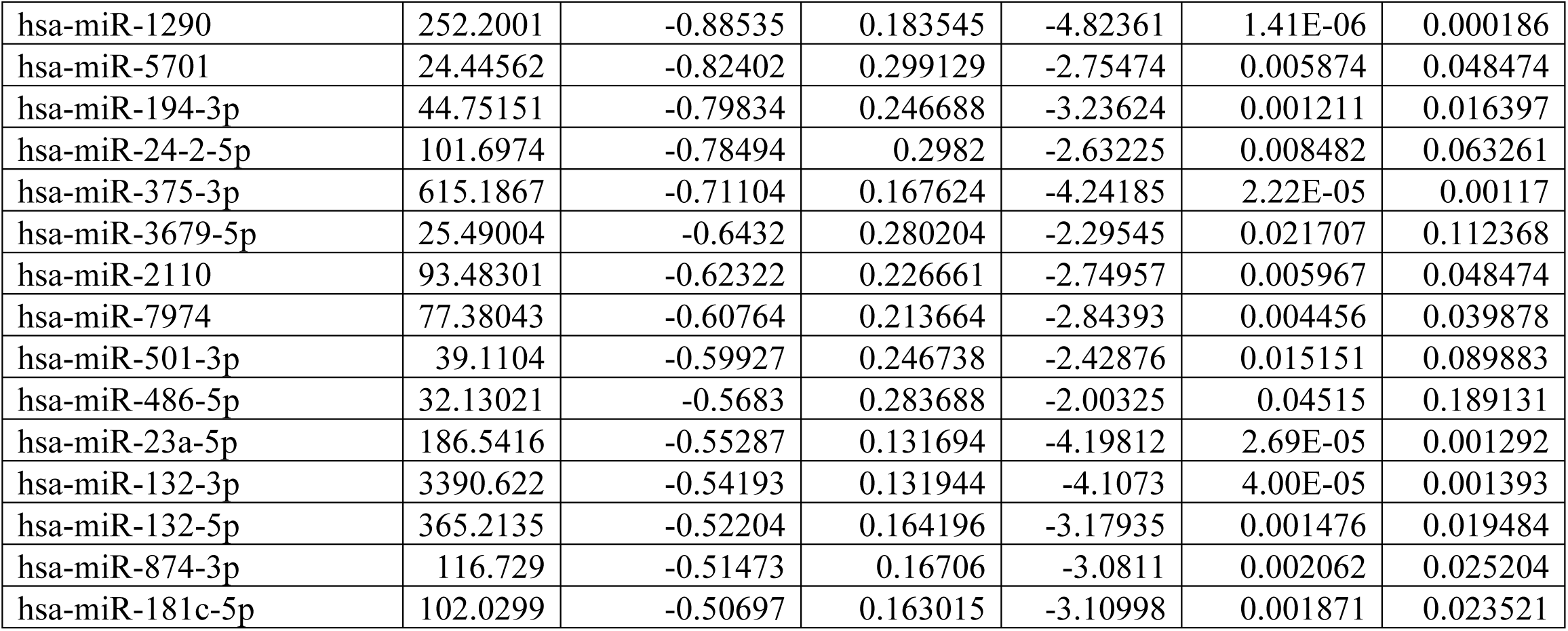
List of P4-DE-miRNAs showing strong upregulation (unadjusted p < 0.05, log2FC > 0.5) and downregulation (unadjusted p < 0.05, log2FC < -0.5) in VK2 cells.

### 3.2 Validation of miRNA sequencing data by quantitative RT-PCR

To validate the microRNA sequencing data, the expression levels of four of the top E2-DE miRNAs (unadjusted p < 0.05 and |log2FC > 0.5|)-hsa-miR-548am-5p, hsa-miR-151b, hsa-miR-181d-3p and hsa-miR-1290 and four of the top P4-DE-miRNAs (unadjusted p < 0.05 and |log2FC > 0.5|)-hsa-miR-182-3p, hsa-miR-582-3p, hsa-miR-1246 and hsa-miR-1290 were quantified by RT-qPCR (n=3; Figure 2A and B). Specific microRNA primers and their gene globe IDs used to validate DE-miRNAs are listed in supplementary table 1. E2- and P4-DE miRNAs that were upregulated in microRNA sequencing results also showed upregulation by RT-qPCR; and those showing downregulation were likewise downregulated by RT-qPCR (Table 4). These findings validate the microRNA sequencing data.

**Figure 2:**
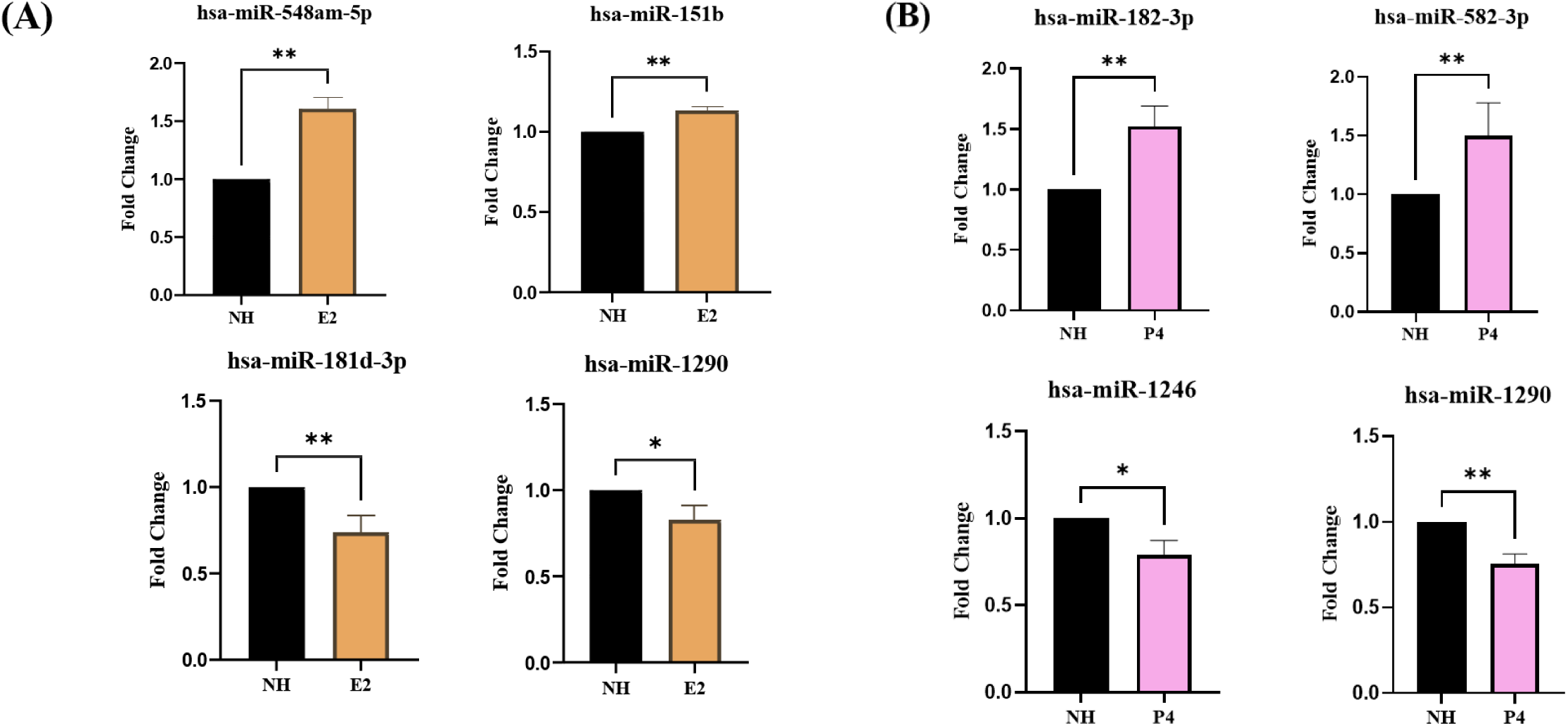
Validation of miRNA sequencing data by RT-qPCR. To validate the miRNA sequencing data, expression levels of (A) top significant differentially regulated E2-miRNAs (unadjusted p < 0.05, |log2FC > 0.5|) and (B) top significant differentially regulated P4-miRNAs (unadjusted p < 0.05, |log2FC > 0.5|) in VK2 cells, compared to NH, were measured by RT-qPCR using hsa-miR-103a-3p as the endogenous control. Data is plotted as mean + SEM of relative fold changes. Samples were run in duplicates, and the data is combined from n=3 independent experiments. An unpaired t-test was used to compare the two treatment groups. Statistical significance is indicated as: ** p < 0.01, * p < 0.05.

**Table 4:**
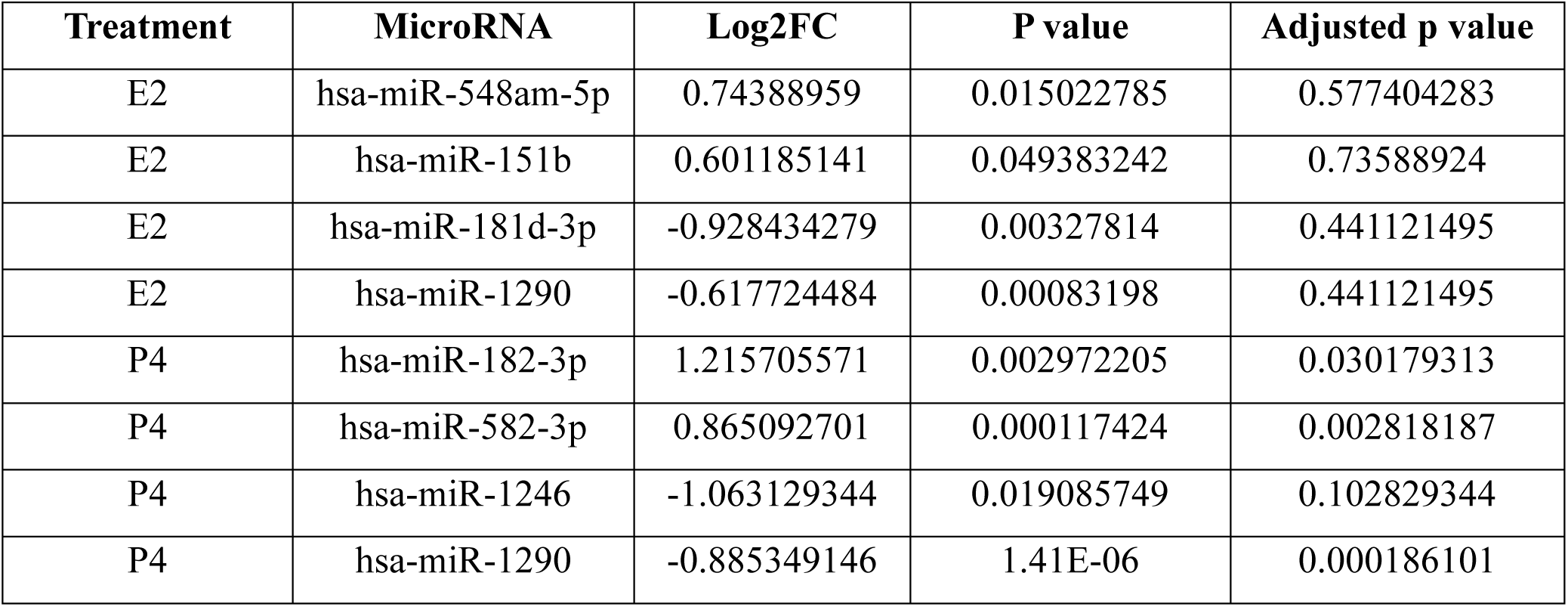
Differentially expressed miRNAs in E2 and P4 treated VK2 cells identified by sequencing (unadjusted p < 0.05).

### 3.3 E2 and P4 have unique microRNA signatures in addition to a set of common microRNAs in VK2 cells

To compare the miRNA profile of E2 and P4 treated VK2 cells, the DE-miRNAs from both treatments (unadjusted p < 0.05) compared to NH, were used to create a Venn diagram. While 13 and 102 miRNAs were regulated by E2 and P4, respectively, 29 miRNAs were regulated by both E2 and P4 (Figure 3A, supplementary table 4). This data indicates that P4 exerts a more pronounced effect on microRNA expression in VK2 cells than E2. Upon visualization of the expression patterns of the 29 common miRNAs, it is evident that many of them (eg, hsa-miR-194-3p, hsa-miR-181c-5p, hsa-miR-210-3p, hsa-miR-3157-5p) are expressed equally under both hormone treatments (Figure 2B). In contrast, some miRNAs (eg, hsa-miR-12136, hsa-miR-4485-3p, hsa-miR-542-3p and hsa-miR-30d-3p) exhibit a higher expression with P4 treatment as compared to E2. None showed higher expression in E2 compared to P4. Interestingly, all 29 common miRNAs show the same directionality of fold change between E2 and P4 but different magnitudes of expression. This suggests that although E2 and P4 both regulate some miRNAs in VK2 cells, the extent of regulation may differ between both hormone treatments.

**Figure 3:**
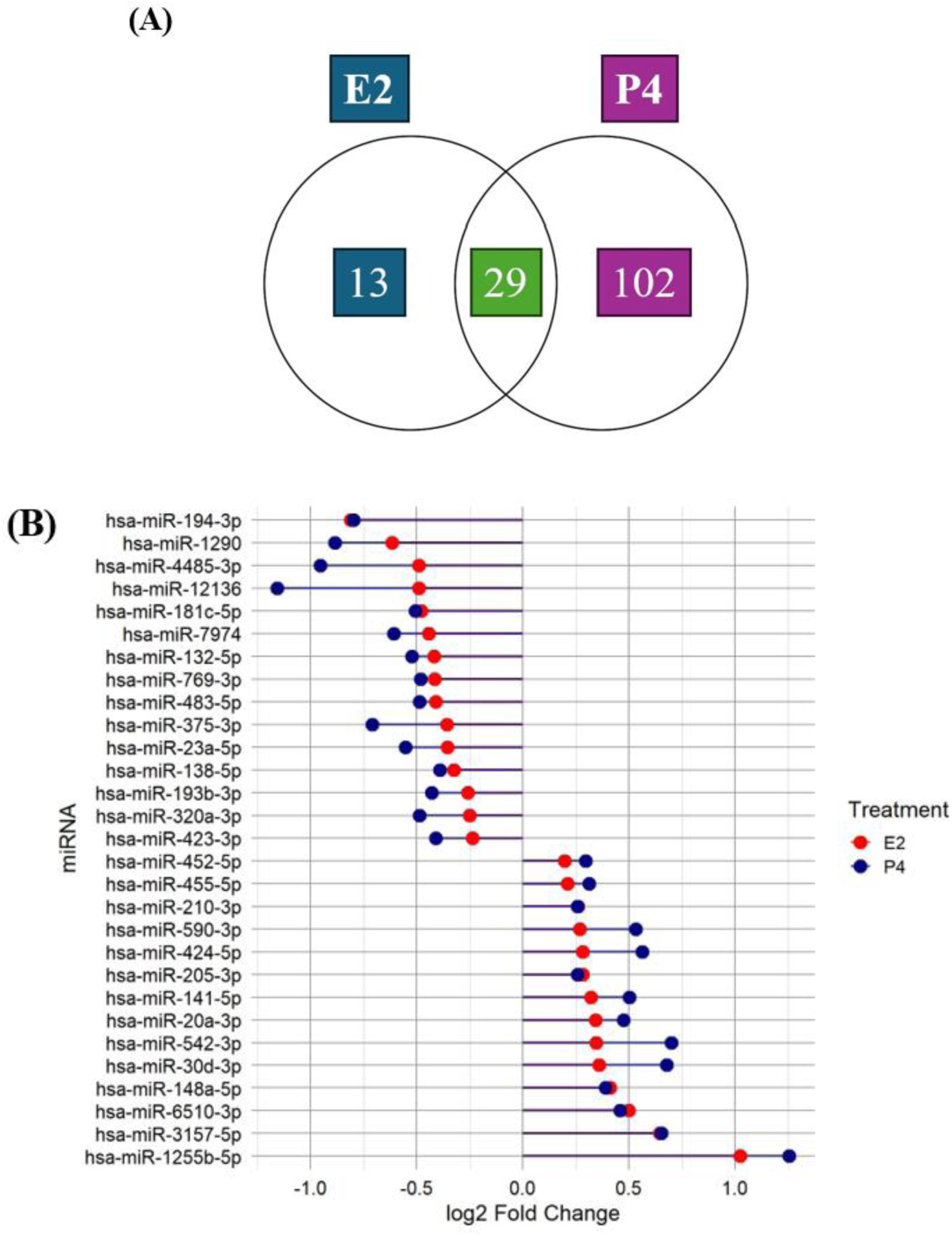
**E2 and P4 have unique microRNA signatures in addition to a set of common microRNAs in VK2 cells. (**A) Venn diagram depicting unique and common significant DE-microRNAs in VK2 cells treated with E2 and P4 (both unadjusted p < 0.05). (B) Lollipop plot depicting the expression pattern of 29 common DE-microRNAs by their log2 fold change values, identified in E2 and P4 treated VK2 cells. Every miRNA is plotted as paired circles to show their differences in magnitudes between both hormones (E2 in red and P4 in blue).

### 3.4 Pathway enrichment analysis of mRNA targets regulated by E2-miRNAs

The sex hormone-mediated miRNA regulation can influence gene expression and signaling pathways in vaginal epithelial cells. To try to understand the biological role of the unique differentially expressed miRNAs in E2 treated VK2 cells (E2-DE-miRNAs) (unadjusted p <0.05), first, mRNA targets of the miRNAs were extracted from miRTaRBase v9.0^[34]^ in miRNet 2.0^[33]^. miRTaRBase only includes experimentally validated miRNA-mRNA targets and excludes the rest, to increase the strength and reliability of the interactions. Figure 4A represents a comprehensive E2-responsive miRNA-mRNA interaction network visualized in miRNet 2.0, highlighting the extensive regulatory breadth, in which every miRNA targets several mRNAs and each mRNA is regulated by multiple miRNAs. Interestingly, four miRNAs with highest number of mRNA targets-hsa-miR-548c-5p (204), hsa-miR-548o-5p, hsa-miR-548au-5p and hsa-miR-548am-5p (203 each) formed a distinct cluster, targeting a shared group of mRNAs in the network, indicating their potentially strong regulatory role (Figure 4A). hsa-miR-99a-5p (136 targets) and hsa-miR-9-3p (109 targets) were two other miRNAs with more than 100 target mRNAs.

**Figure 4:**
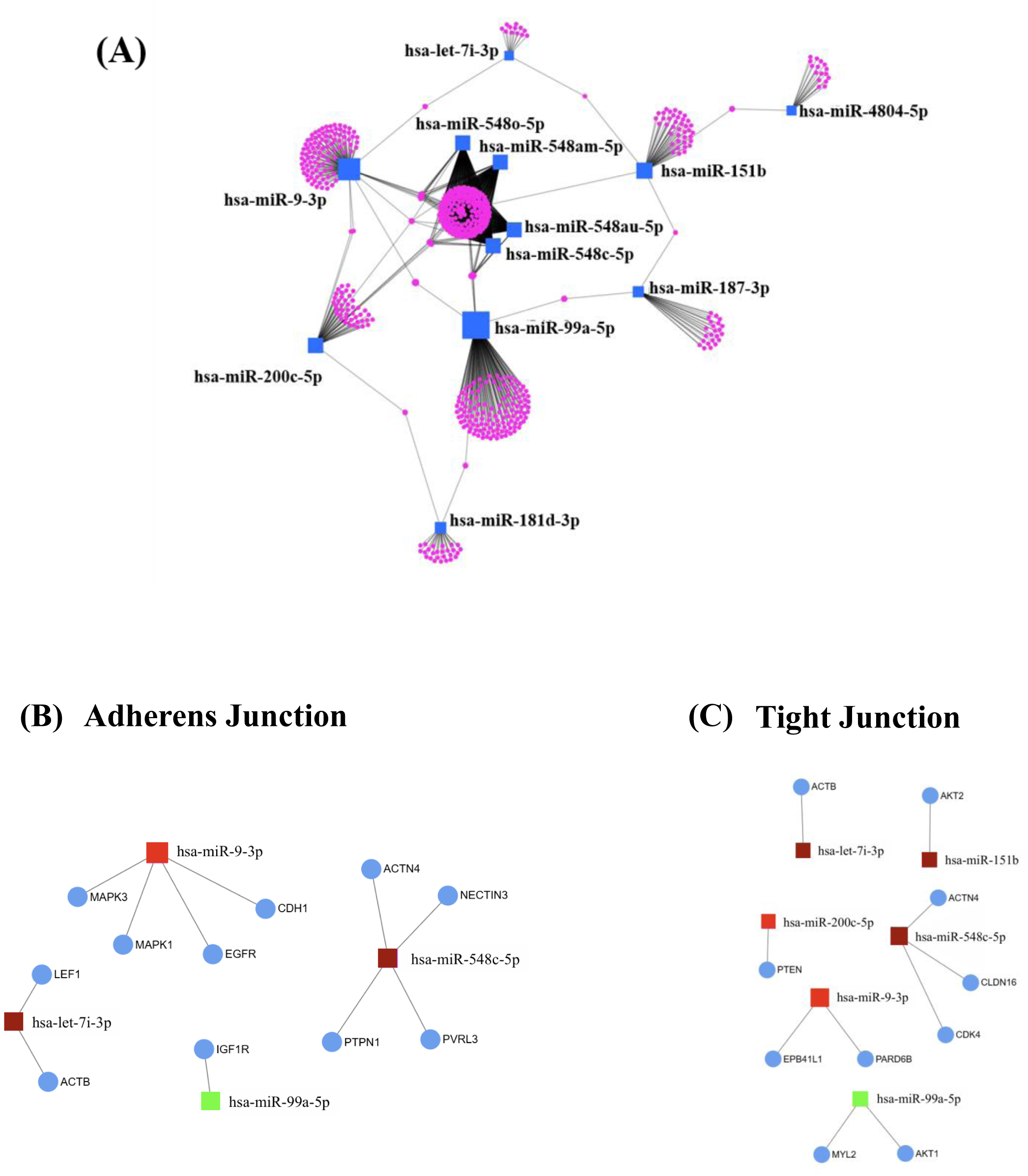
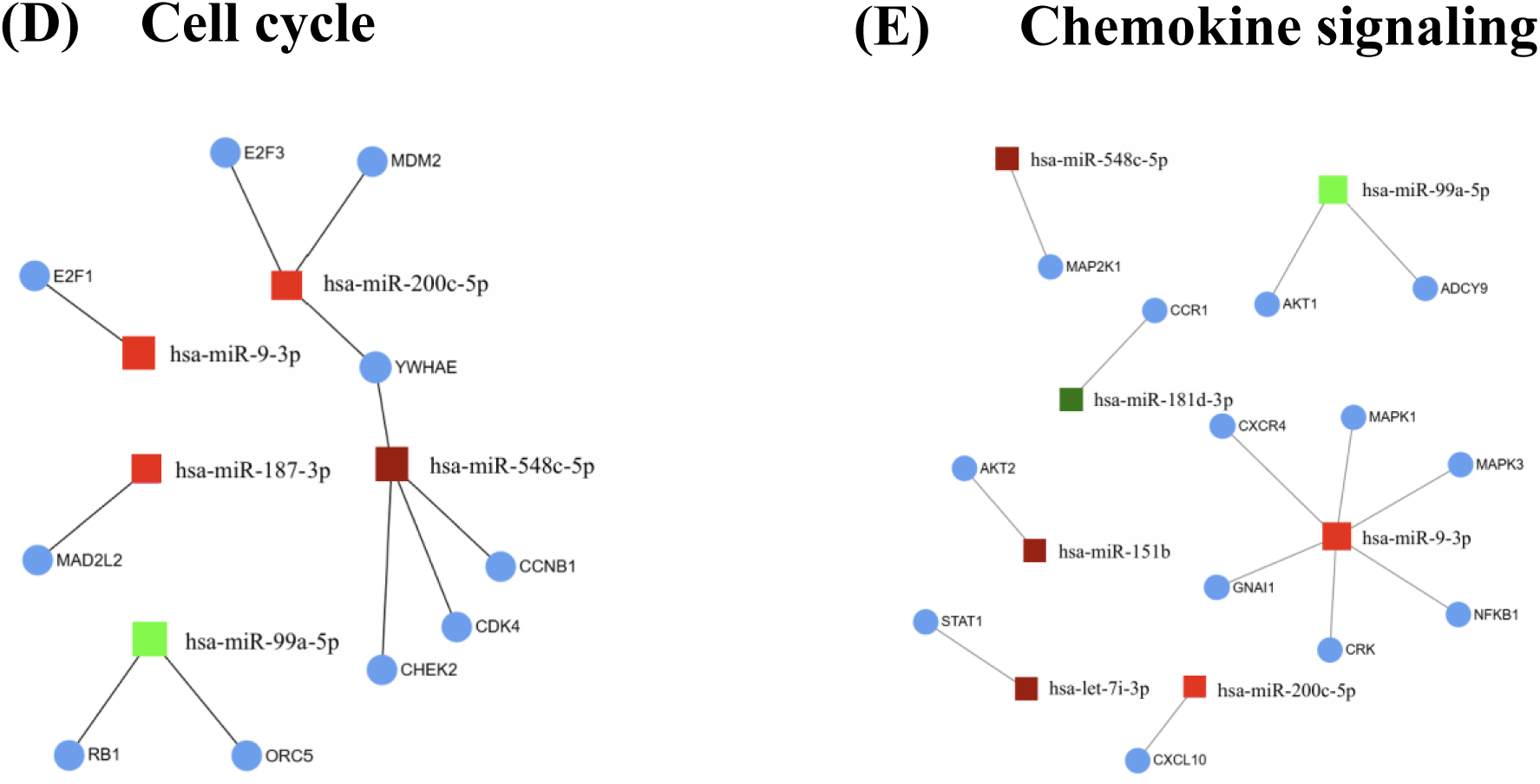
Pathway enrichment analysis of mRNA targets of miRNAs regulated by E2. (A) Interaction network of unique E2-DE-miRNAs (unadjusted p < 0.05) in blue squares and their experimentally validated mRNA targets extracted from miRTaRBase v9 (pink circles). (B-E) KEGG pathway enrichment analysis was performed on the mRNA targets of unique miRNAs regulated by E2. To understand the role of E2-DE-miRNAs in regulating vaginal epithelial cell functions related to viral susceptibility, significant pathways (FDR < 0.05) regulated by 10 or more mRNA targets, associated with epithelial barrier integrity, cell development and innate immune response were used to create specific pathway sub-networks. Within every sub-network, dark-red squares represent miRNAs with strong upregulation (log2FC > 0.5), and light-red colored squares represent miRNAs with moderate upregulation (0 < log2FC < 0.5). The dark green squares represent miRNAs with strong downregulation (log2FC < -0.5) and light-green squares represent miRNAs with moderate downregulation (0 > log2FC > -0.5). The target mRNAs are depicted in blue circles.

Next, to examine the signaling pathways regulated by mRNA targets of unique E2-DE-miRNAs, a KEGG pathway enrichment was performed on the mRNA targets in the network using hypergeometric overrepresentation approach. It yielded 34 significant pathways (FDR < 0.05) (see methods 2.6; Table 5) including glioma and HTLV-1 infection (14 mRNA hits), insulin signaling pathway (13 mRNA hits), oocyte meiosis (12 mRNA hits), progesterone mediated oocyte maturation (11 mRNA hits), mTOR signaling (9 mRNA hits), apoptosis (8 mRNA hits), endometrial cancer (7 hits) among others. To understand the role of E2-DE-miRNAs in regulating vaginal epithelial cell functions related to viral susceptibility, subnetworks of pathways involving 10 or more target mRNAs, associated with epithelial barrier integrity (adherens junction-11 mRNA hits, tight junction-10 mRNA hits); cell growth and development (cell cycle -10 mRNA hits); and innate immune responses (chemokine signaling pathway-13 mRNA hits) were generated (Figure 4B-E). These DE-miRNAs in VK2 cells could possibly be the key mediators of E2 associated enhanced protection against STIs.

**Table 5:**
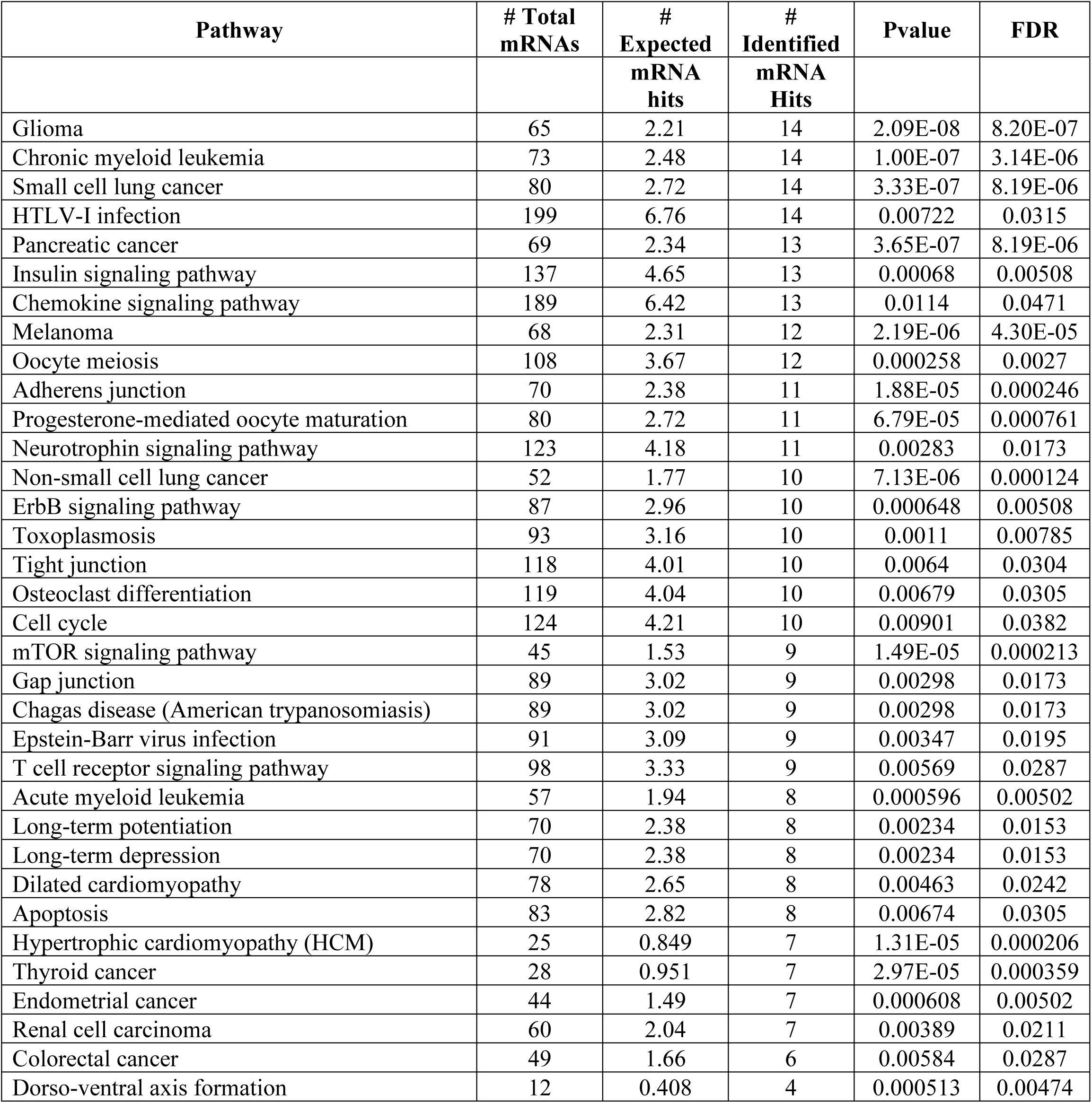
KEGG Pathway enrichment analysis of mRNA targets of unique E2-DE-miRNAs in VK2 cells (FDR < 0.05). Significant pathways overlapping with those enriched by mRNA targets of 20 random miRNAs were removed.

### 3.5 Pathway enrichment analysis of mRNA targets regulated by P4-miRNAs

Next, to elucidate the role of unique miRNAs regulated by P4 (P4-DE-miRNAs; unadjusted p < 0.05) in influencing signaling pathways in VK2 cells, the mRNA targets of P4-DE-miRNAs were extracted from miRTaRBase v9.0, and a comprehensive P4-responsive miRNA-mRNA network was visualized in miRNet 2.0. Because of the large size of the network containing 9240 mRNA targets, a degree filter of 10 was applied to get a condensed, less complex interaction network (Figure 5A). hsa-miR-20a-5p and hsa-miR-106b-5p were the largest nodes identified in the network targeting 105 and 104 mRNAs, respectively, indicating the breadth of their regulatory function.

**Figure 5:**
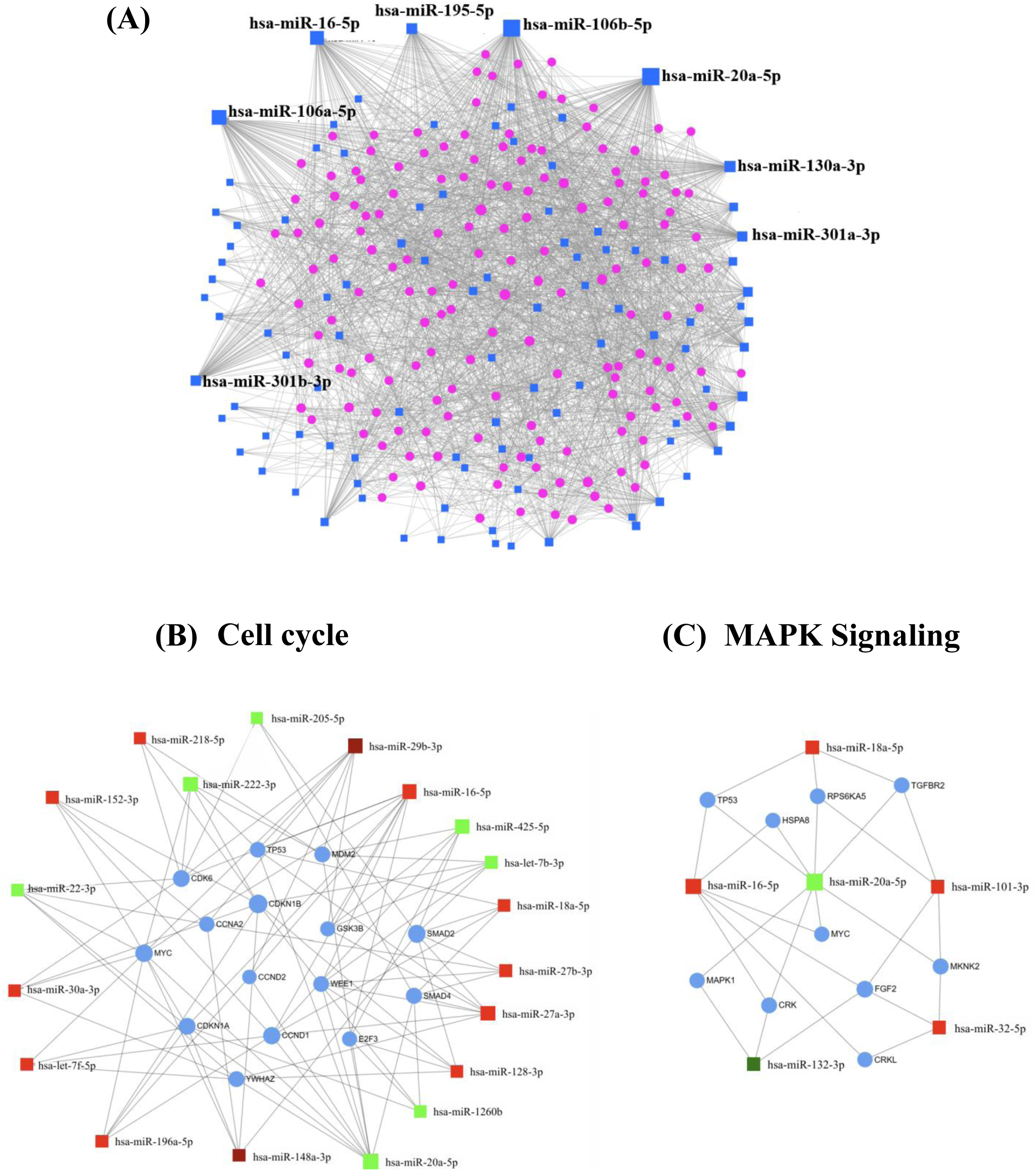
Pathway enrichment analysis of mRNA targets of miRNAs regulated by P4. (A) Interaction network of unique P4-DE-miRNAs (unadjusted p < 0.05, degree filter 10) in blue squares and their experimentally validated mRNA targets extracted from miRTaRBase v9 (pink circles). (B and C) KEGG pathway enrichment analysis was performed on the mRNA targets of unique P4-regulated miRNAs. To understand the role of P4-DE-miRNAs in regulating vaginal epithelial cell functions related to viral susceptibility, significant pathways (FDR < 0.05) regulated by 10 or more mRNA targets, associated with cell development and innate immune response were used to create sub-networks. No significant pathways related to epithelial barrier integrity with 10 or more target mRNAs were identified. Within every sub-network, dark-red squares represent miRNAs with strong upregulation (log2FC > 0.5), and light-red colored squares represent miRNAs with moderate upregulation (0 < log2FC < 0.5). The dark green squares represent miRNAs with strong downregulation (log2FC < -0.5) and light-green squares represent miRNAs with moderate downregulation (0 > log2FC > -0.5). The target mRNAs are depicted in blue circles.

KEGG pathway enrichment of all mRNA targets within the network identified 33 significant pathways (FDR < 0.05) (Table 6), including cell cycle (15 mRNA hits), glioma (10 mRNA hits), MAPK signaling (10 mRNA hits), p53 signaling (9 mRNA hits), Wnt signaling (8 mRNA hits), adherens junction (7 mRNA hits), JAK-STAT signaling (6 mRNA hits), endocytosis (5 mRNA hits) and several others. To understand the role of P4-DE-miRNAs in regulating vaginal epithelial cell functions related to viral susceptibility, subnetworks of pathways involving 10 or more mRNA targets; associated with cell growth and development (cell cycle-15 mRNA hits) and innate immune responses (MAPK signaling-10 mRNA hits), were generated (Figure 5 B and C). No pathway related to epithelial barrier integrity, regulated by 10 or more target mRNAs was identified. Although pathways such as cell cycle, chemokine signaling, apoptosis and others were also found to be regulated by mRNA targets of E2-DE-miRNAs, the microRNAs involved were different than P4. This data suggests that multiple DE-miRNAs can regulate a biological process to achieve similar or opposite functional outcomes.

**Table 6:**
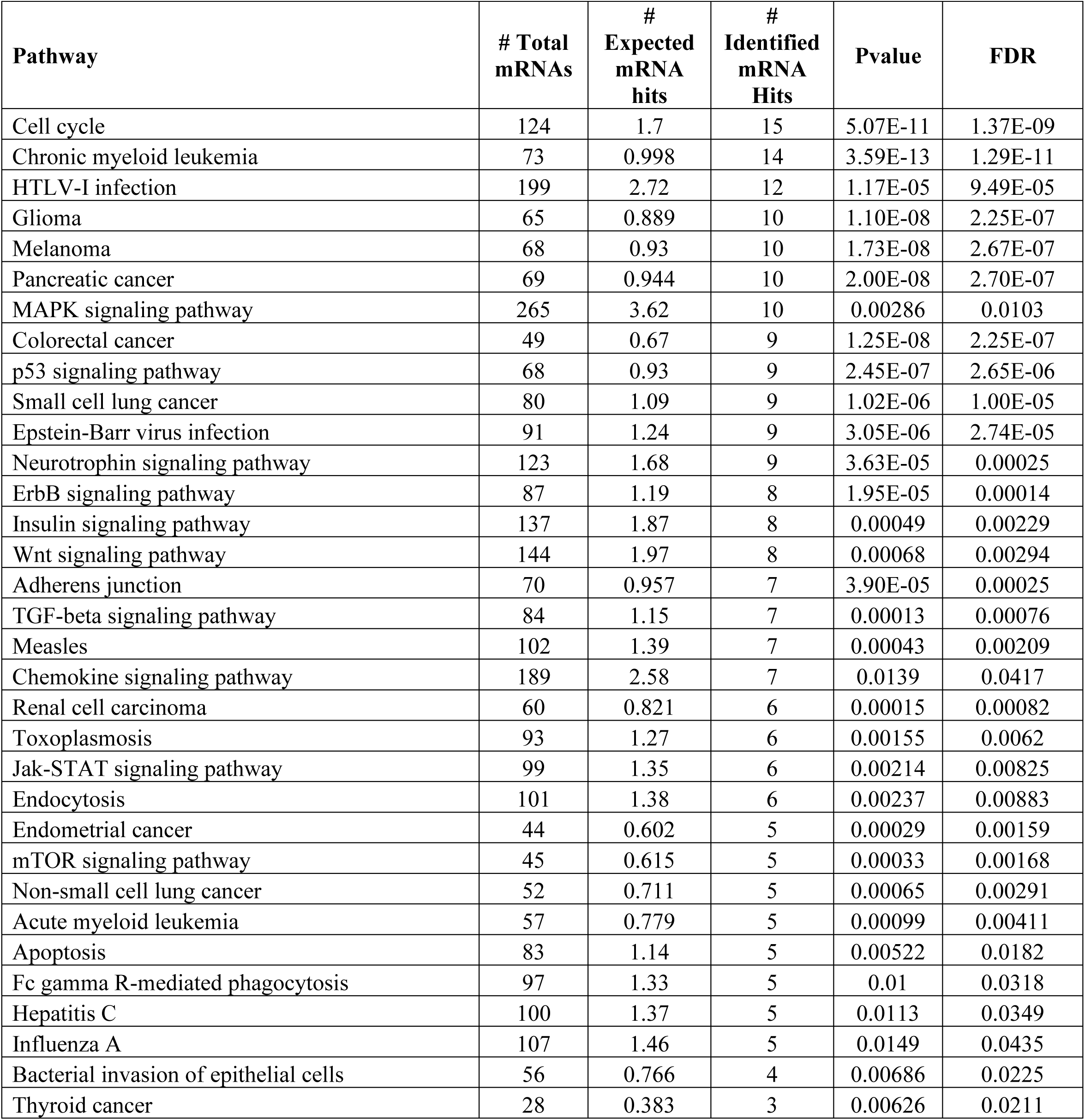
KEGG Pathway enrichment analysis of mRNA targets of unique P4-DE-miRNAs in VK2 cells (FDR < 0.05). Significant pathways overlapping with those enriched by mRNA targets of 20 random miRNAs were removed.

## Discussion

While microRNA research is advancing our understanding of the importance of miRNAs in regulating host biological processes, influencing disease outcomes and developing novel therapeutic strategies^[35]^, no studies in vaginal epithelial cells have examined the regulation of miRNA expression by sex hormones. Here, we show for the first time the effect of physiological concentrations of E2 or P4 on microRNA expression in vaginal epithelial cells (VK2 cells) that regulate post-transcriptional gene expression. In this study, we demonstrated that E2 and P4 treated VK2 cells have unique microRNA signatures in addition to a few miRNAs which were regulated by both E2 and P4, highlighting the regulation of these microRNAs by sex hormones. Furthermore, we did an analysis to understand the functional role of significant DE-miRNAs in VK2 cells after hormone treatment. Overall, the 13 unique miRNAs regulated by E2 and 102 unique miRNAs regulated by P4 were identified in VK2 cells. Most of the pathways regulated by the mRNA targets of E2 and P4-DE-miRNAs are related to cell-cell adhesion, cell growth and development and innate immune responses; all of which influence epithelial barrier integrity and susceptibility to sexually transmitted pathogens^[3]^. We also validated the top E2 and P4-DE-miRNAs identified in the transcription analysis by quantifying the miRNA expression in VK2 cells, by RT-qPCR. Together, this data offers valuable mechanistic insights into the fine regulation by sex hormone driven miRNAs in vaginal epithelial cells, potentially influencing susceptibility to sexually transmitted pathogens in women.

Our work has demonstrated that in VK2 cells, physiological levels of E2, regulate miRNAs whose mRNA targets are involved in influencing adherens junction and tight junction pathways. This is consistent with previous studies demonstrating E2-mediated enhancement of epithelial barrier integrity in the lower FRT^[14, 36, 37]^. We observed an upregulation of hsa-miR-9-3p in VK2 cells potentially targeting PARD6B mRNA, a homolog of PARD6, in the tight junction pathway (Figure 4C). It acts as a scaffold protein in the Par complex that plays a crucial role in regulation of cell polarity and tight junction assembly^[38, 39]^. Cunliffe et al. observed that inhibition of PARD6B in MCF7 breast cancer cells led to disruption of tight junction networks^[39]^. However, they found enhanced levels of cytoplasmic ZO-1, suggesting that loss of PARD6B could potentially affect sub-cellular localization of ZO-1 but may not reduce overall protein expression. The same study also reported that inhibition of PARD6B did not disrupt assembly of adherens junctions in MCF7 cells^[39]^. Another potential mRNA target of hsa-miR-9-3p identified in our analysis was E-cadherin (CDH1), a key component of adherens junctions. (Figure 4B). This is supported by Sui et al. who observed upregulation of hsa-miR-9-3p in primary ovarian cancer tissues that promoted epithelial-mesenchymal transition (EMT) by targeting E-cadherin (CDH1), leading to the disruption of adherens junctions^[40]^. Currently, there are no studies that directly link hsa-miR-9-3p to epithelial barrier integrity and existing literature focuses on its role in regulating EMT process in cancer. Based on our analysis, hsa-miR-9-3p represents one potential regulator of mRNAs in cell junction pathways, under the influence of E2. However, several other E2-DE-miRNAs were also identified whose mRNA targets were a part of adherens and tight junction pathways in VK2 cells. Since these pathways are highly interconnected, the combined function of E2-miRNAs may strengthen overall epithelial barrier integrity to prevent pathogen entry in vaginal tract. However, this requires experimental validation.

Cell proliferation is a hallmark of vaginal epithelium and the proliferative effects of E2 on epithelial cells of vaginal tract are well established^[41]^. Our work identified mRNA targets of several E2-DE-miRNAs to be involved in cell cycle pathway in VK2 cells. Interestingly, YWHAE appeared to be a potential target of both hsa-miR-200c-5p and hsa-miR-548c-5p (Figure 4D). These are tumor suppressor miRNAs and are crucial in preventing uncontrolled cell proliferation. YWHAE plays a significant role in regulating multiple biological processes such as cell proliferation, apoptosis and signal transduction and a negative regulator of CDC25 phosphatases that regulate cell cycle progression ^[42]^. Although YWHAE has been associated with excessive cell proliferation and invasion in several cancers^[42–44]^, our data indicates YWAHE is likely to be downregulated by E2-miRNAs, consistent with controlled proliferation of vaginal epithelial cells, rather than malignant growth. In our analysis, hsa-miR-548c-5p was identified as a potential regulator of CCNB1, a key gene involved in mitosis, and overexpression of CCNB1 has been shown to promote tumor growth^[45]^. Together, our data indicates miRNA-mediated regulation of mRNAs that are part of the cell cycle pathway under the influence of E2, however, further experimental evaluation is necessary to test whether the pathway is activated or inhibited.

The immune responses in the FRT are tightly regulated by sex hormones and consistent with the immunosuppressive role of E2, we observed enrichment of chemokine signaling pathway regulated by mRNA targets of E2-DE-miRNAs^[1]^. Our analysis identified multiple pro-inflammatory genes NFKB1, MAPK1 and MAPK3, as potential targets of hsa-miR-9-3p.. NF-κB1 (p50) is a member of NF-κB family, that regulates production of pro-inflammatory cytokines and chemokines^[46]^. Interestingly, Prezioso et al., observed downregulation of NF-κB and reduced production of pro-inflammatory cytokines-IL-6, IL-1β, and TNFα, associated with miR-9 expression in COVID-19 patients; highlighting its anti-inflammatory role ^[47]^. MAPK signaling plays a crucial role in recruitment of immune cells and regulating proinflammatory gene expression^[48]^. We also identified MAPK1 (ERK2) and MAPK3 (ERK1), key mediators of MAPK signaling pathway, as potential targets of hsa-miR-9-3p. Additionally, MAP2K1 (MEK1), acting upstream of the ERKs in MAPK signaling was found to be a potential target of hsa-miR-548c-5p- known negative regulator of MAP2K1. ^[49]^. Since both MAPK and NF-κB signaling function parallelly and regulate the expression of pro-inflammatory cytokines and chemokines, it is tempting to speculate that E2-mediated dampening of excessive inflammation in VK2 cells could be regulated by mRNA targets of E2-DE-miRNAs, however, this needs to be verified experimentally.

Our analysis of the mRNA targets of unique P4-DE-miRNAs showed enrichment of the cell cycle pathway in vaginal epithelial cells. Although this pathway was also enriched by the mRNA targets of E2-DE-miRNAs, the specific miRNAs involved were unique to E2. Previous studies show that P4 inhibits E2-induced cell proliferation and promotes maturation of epithelial cells in the FRT^[50, 51]^. In line with this, we identified potential mRNA targets of multiple differentially expressed miRNAs in P4 treated VK2 cells including regulators of cell cycle, such as CDKN1A, CCND1, MYC, TP53, MDM2, and E2F3 (Figure 5B). Notably, CDKN1A (p21), a key inhibitor of cell cycle progression could likely be targeted by upregulated hsa-let-7f-5p and downregulated hsa-miR-20a-5p in VK2 cells. hsa-let-7 family is a tumor suppressor, targeting oncogenes and regulators of cell cycle progression including CDKN1A, CCND1 and MYC, thereby arresting the cells at G1/S checkpoint (reviewed in Wang et al., 2024)^[52]^. Previously, using transcriptomic profiling, we determined that treatment of vaginal epithelial cells with medroxyprogesterone acetate, a hormonal contraceptive that mimics the effects of progesterone, resulted in downregulation of genes required for cell cycle and cell division^[5]^. hsa-miR-20a-5p, a member of miR-17/92 cluster is known to promote cell cycle progression by repressing CDKN1A^[53]^. But, in our data, the downregulation of has-miR-20a-5p could potentially upregulate CDKN1A, leading to the inhibition of cell cycle. Other differentially expressed miRNAs such as hsa-miR-16-5p and hsa-miR-27a-3p are also known to inhibit cell cycle^[54, 55]^, however, the combined action of all miRNAs and their target mRNAs will determine activation or inhibition of the pathway. Although these findings raise the possibility that P4 may negatively affect cell cycle progression through microRNA regulation, functional validation is required to confirm this mechanism.

Additionally, mRNA targets of P4-DE-miRNAs were involved in MAPK signaling pathway in VK2 cells (Figure 5C). In contrast to E2, where MAPK1 was identified as a potential target of upregulated has-miR-9-3p (likely reducing MAPK1 expression), opposite trends were detected in P4-treated VK2 cells. MAPK1 appeared to be targeted by hsa-miR-132-3p and hsa-miR-20a-5p, two downregulated miRNAs, likely enhancing MAPK1 expression, and the production of pro-inflammatory cytokines and chemokines via AP1-dependent transcription^[48]^. Our findings are supported by He et al. who reported upregulated MAPK1 with reduced expression of hsa-miR-132-3p in chronic fluorosis model of SH-SY5Y neuronal cells^[56]^. In addition, TP53 was a potential target of three miRNAs-upregulated has-miR-18a-5p and hsa-miR-16-5p, and downregulated hsa-miR-20a-5p. Given TP53’s role in regulating inflammatory response, functional studies will be needed to determine which miRNA has a more dominant effect on its expression^[57]^. Together, these results highlight that P4-mediated enhancement of inflammatory responses in vaginal epithelial cells could potentially be regulated by P4-DE-miRNAs.

This study is the first to investigate the effect of female sex hormones on miRNA expression in vaginal epithelial cells grown in ALI cultures, *in vitro*, but there are some limitations. We characterized the vaginal epithelial cell miRNA profile in the absence of stromal cells which are known to interact with the former in the FRT and could potentially influence their miRNA landscape^[50, 58]^. The findings of this study need to be validated using primary vaginal epithelial cells isolated from whole vaginal tissues. MicroRNA function is cell type specific, and even though we discussed the role of only a few DE-miRNAs and their potential mRNA targets in regulating each pathway, several miRNAs were identified in both hormone treatments. Co-ordination of multiple miRNAs is necessary to regulate specific biological functions, reflecting the complexity of miRNA regulation^[17]^. Given the tendency of identifying non-specific signaling pathways in enrichment analysis, we performed a KEGG pathway enrichment on target mRNAs of 20 random miRNAs and eliminated pathways that were significant (FDR < 0.05) from our E2 and P4 results. Although important pathways related to epithelial barrier function, cell development and immune responses were not significantly enriched by the mRNA targets of these random miRNAs (Table 1), many were close to the significance threshold of FDR < 0.05. This emphasizes the need for functional validation using miRNA mimics and inhibitors to establish the causal roles of E2- and P4-DE-miRNAs, and their mRNA targets in regulating signaling pathways.

In conclusion, we have demonstrated that physiological concentrations of female sex hormones E2 and P4, regulate microRNA expression in vaginal epithelial cells, influencing mRNAs that are associated with epithelial cell junctions, cell cycle and innate immune signaling pathways. This study provides a deeper insight into hormone-specific microRNA signatures in the vaginal epithelial cells, affecting epithelial barrier function and susceptibility to STIs in women, making them potential targets for therapeutic interventions.

## Acknowledgements

This study was supported by research grant # 159229 from Canadian Institutes of Health Research (CIHR) to Dr. Charu Kaushic.

## Ethics statement

The authors confirm that no ethical approval was required as this study was performed using a cell line and not primary cells or *in-vivo* models.

## Author Contribution Statement

Shreya Joshi: Conceptualization, Investigation, Data analysis, Writing-original draft, review and editing. Aisha Nazli: Investigation, Writing-review and editing. Chris Verschoor: Data analysis, Writing-review and editing. Charu Kaushic: Conceptualization, Funding acquisition, Project management, Supervision, Writing-review and editing.

