## Supplementary Tables for "Transcriptomic profiling reveals estradiol and progesterone specific microRNA signatures differentially regulate mRNA targets in vaginal epithelial cells"

### Supplementary Data

**Supplementary table 1**: microRNA primers used in RT-qPCR analysis and their Gene Globe IDs.

| microRNA | Qiagen Gene Globe ID |
|---|---|
| hsa-miR-548am-5p | YP00205882 |
| hsa-miR-151b | YP02110935 |
| hsa-miR-181d-3p | YP02105350 |
| hsa-miR-1290 | YP02118634 |
| hsa-miR-182-3p | YP00204098 |
| hsa-miR-582-3p | YP00204072 |
| hsa-miR-1246 | YP00205630 |
| hsa-miR-103a-3p | YP00204063 |

**Supplementary table 2:** Complete list of differentially expressed miRNAs in E2 treated VK2 cells (unadjusted $p < 0.05$).

| miRNA | base mean | Log2 Fold Change (FC) | lfc SE | stat | pvalue | padj |
|---|---|---|---|---|---|---|
| hsa-miR-181d-3p | 20.69790527 | -0.92843 | 0.315754 | -2.94038 | 0.003278 | 0.441121 |
| hsa-miR-194-3p | 44.7515108 | -0.81245 | 0.255309 | -3.18222 | 0.001462 | 0.441121 |
| hsa-miR-4804-5p | 23.59773089 | -0.6616 | 0.30109 | -2.19736 | 0.027995 | 0.577404 |
| hsa-miR-1290 | 252.2001009 | -0.61772 | 0.184841 | -3.34193 | 0.000832 | 0.441121 |
| hsa-miR-4485-3p | 385.0194657 | -0.4899 | 0.174366 | -2.80963 | 0.00496 | 0.441121 |
| hsa-miR-12136 | 7860.422702 | -0.48873 | 0.214205 | -2.28158 | 0.022514 | 0.577404 |
| hsa-miR-181c-5p | 102.0298855 | -0.4798 | 0.167804 | -2.85927 | 0.004246 | 0.441121 |
| hsa-miR-7974 | 77.38042972 | -0.44268 | 0.217446 | -2.03583 | 0.041767 | 0.705524 |
| hsa-miR-132-5p | 365.2135438 | -0.41737 | 0.165609 | -2.52021 | 0.011728 | 0.523593 |
| hsa-miR-769-3p | 98.42823591 | -0.41642 | 0.164758 | -2.52748 | 0.011489 | 0.523593 |
| hsa-miR-483-5p | 170.7346123 | -0.40898 | 0.141191 | -2.89667 | 0.003771 | 0.441121 |
| hsa-miR-375-3p | 615.1866525 | -0.35855 | 0.168071 | -2.1333 | 0.0329 | 0.604776 |

| | | | | | | |
|---|---|---|---|---|---|---|
| hsa-miR-23a-5p | 186.5416015 | -0.35609 | 0.133943 | -2.65853 | 0.007848 | 0.520836 |
| hsa-miR-138-5p | 3071.165489 | -0.32432 | 0.125909 | -2.57583 | 0.01 | 0.520836 |
| hsa-miR-99a-5p | 811.2803729 | -0.26581 | 0.120575 | -2.20452 | 0.027487 | 0.577404 |
| hsa-miR-193b-3p | 442.8027388 | -0.25994 | 0.132314 | -1.96453 | 0.049468 | 0.735889 |
| hsa-miR-320a-3p | 6053.098655 | -0.25037 | 0.104715 | -2.39101 | 0.016802 | 0.577404 |
| hsa-miR-423-3p | 4044.026906 | -0.23748 | 0.103532 | -2.29383 | 0.0218 | 0.577404 |
| hsa-miR-9-3p | 1314.673269 | 0.193553 | 0.087571 | 2.210248 | 0.027088 | 0.577404 |
| hsa-miR-452-5p | 1138.977807 | 0.195989 | 0.083516 | 2.346737 | 0.018939 | 0.577404 |
| hsa-miR-455-5p | 2449.418747 | 0.21068 | 0.095401 | 2.208362 | 0.027219 | 0.577404 |
| hsa-miR-200c-5p | 521.0404867 | 0.220749 | 0.10963 | 2.013573 | 0.044054 | 0.706 |
| hsa-miR-210-3p | 3664.649761 | 0.261436 | 0.120005 | 2.178534 | 0.029366 | 0.577404 |
| hsa-miR-590-3p | 715.6764482 | 0.267699 | 0.11519 | 2.323984 | 0.020126 | 0.577404 |
| hsa-miR-424-5p | 3183.01585 | 0.281441 | 0.136971 | 2.054746 | 0.039904 | 0.69277 |
| hsa-miR-205-3p | 1233.261527 | 0.282419 | 0.129094 | 2.187698 | 0.028692 | 0.577404 |
| hsa-miR-141-5p | 847.642559 | 0.320964 | 0.11424 | 2.809566 | 0.004961 | 0.441121 |
| hsa-miR-20a-3p | 157.0335132 | 0.34352 | 0.170416 | 2.01578 | 0.043823 | 0.706 |
| hsa-miR-542-3p | 534.0271877 | 0.345195 | 0.133451 | 2.586675 | 0.009691 | 0.520836 |
| hsa-miR-30d-3p | 94.78351141 | 0.359013 | 0.169291 | 2.120683 | 0.033948 | 0.606223 |
| hsa-miR-148a-5p | 101.257424 | 0.411543 | 0.185324 | 2.220661 | 0.026374 | 0.577404 |
| hsa-miR-187-3p | 60.43332044 | 0.464618 | 0.202181 | 2.298036 | 0.02156 | 0.577404 |
| hsa-miR-6510-3p | 110.8952514 | 0.500404 | 0.180805 | 2.767641 | 0.005646 | 0.441121 |
| hsa-let-7i-3p | 81.94956987 | 0.529482 | 0.244711 | 2.163706 | 0.030487 | 0.577404 |
| hsa-miR-151b | 33.44182486 | 0.601185 | 0.305905 | 1.965268 | 0.049383 | 0.735889 |
| hsa-miR-3157-5p | 30.58668837 | 0.646263 | 0.267895 | 2.412379 | 0.015849 | 0.577404 |
| hsa-miR-548c-5p | 33.74432544 | 0.657464 | 0.279444 | 2.352757 | 0.018635 | 0.577404 |
| hsa-miR-548o-5p | 33.74432544 | 0.657464 | 0.279444 | 2.352757 | 0.018635 | 0.577404 |
| hsa-miR-548am-5p | 28.77837032 | 0.74389 | 0.305897 | 2.431829 | 0.015023 | 0.577404 |
| hsa-miR-101-2-5p | 9.046825449 | 1.001797 | 0.510019 | 1.964235 | 0.049503 | 0.735889 |
| hsa-miR-1255b-5p | 11.90627073 | 1.024314 | 0.472485 | 2.16793 | 0.030164 | 0.577404 |

| | | | | | | |
|---|---|---|---|---|---|---|
| hsa-miR-548au-5p | 13.94809522 | 1.186348 | 0.459004 | 2.584612 | 0.009749 | 0.520836 |

**Supplementary table 3:** Complete list of differentially expressed miRNAs in P4 treated VK2 cells (unadjusted $p < 0.05$).

| **miRNA** | **base mean** | **Log2 Fold Change (FC)** | **lfc SE** | **stat** | **pvalue** | **padj** |
|---|---|---|---|---|---|---|
| hsa-miR-10401-3p | 20.0685 | -1.16046 | 0.333418 | -3.48049 | 0.0005 | 0.008714 |
| hsa-miR-12136 | 7860.423 | -1.15758 | 0.214209 | -5.40398 | 6.52E-08 | 1.72E-05 |
| hsa-miR-1246 | 776.6227 | -1.06313 | 0.453582 | -2.34385 | 0.019086 | 0.102829 |
| hsa-miR-6883-3p | 8.345756 | -1.04369 | 0.496362 | -2.10269 | 0.035493 | NA |
| hsa-miR-4485-3p | 385.0195 | -0.95472 | 0.173987 | -5.48733 | 4.08E-08 | 1.72E-05 |
| hsa-miR-1290 | 252.2001 | -0.88535 | 0.183545 | -4.82361 | 1.41E-06 | 0.000186 |
| hsa-miR-5701 | 24.44562 | -0.82402 | 0.299129 | -2.75474 | 0.005874 | 0.048474 |
| hsa-miR-194-3p | 44.75151 | -0.79834 | 0.246688 | -3.23624 | 0.001211 | 0.016397 |
| hsa-miR-24-2-5p | 101.6974 | -0.78494 | 0.2982 | -2.63225 | 0.008482 | 0.063261 |
| hsa-miR-375-3p | 615.1867 | -0.71104 | 0.167624 | -4.24185 | 2.22E-05 | 0.00117 |
| hsa-miR-3679-5p | 25.49004 | -0.6432 | 0.280204 | -2.29545 | 0.021707 | 0.112368 |
| hsa-miR-2110 | 93.48301 | -0.62322 | 0.226661 | -2.74957 | 0.005967 | 0.048474 |
| hsa-miR-7974 | 77.38043 | -0.60764 | 0.213664 | -2.84393 | 0.004456 | 0.039878 |
| hsa-miR-501-3p | 39.1104 | -0.59927 | 0.246738 | -2.42876 | 0.015151 | 0.089883 |
| hsa-miR-486-5p | 32.13021 | -0.5683 | 0.283688 | -2.00325 | 0.04515 | 0.189131 |
| hsa-miR-23a-5p | 186.5416 | -0.55287 | 0.131694 | -4.19812 | 2.69E-05 | 0.001292 |
| hsa-miR-132-3p | 3390.622 | -0.54193 | 0.131944 | -4.1073 | 4.00E-05 | 0.001393 |
| hsa-miR-132-5p | 365.2135 | -0.52204 | 0.164196 | -3.17935 | 0.001476 | 0.019484 |
| hsa-miR-874-3p | 116.729 | -0.51473 | 0.16706 | -3.0811 | 0.002062 | 0.025204 |
| hsa-miR-181c-5p | 102.0299 | -0.50697 | 0.163015 | -3.10998 | 0.001871 | 0.023521 |
| hsa-miR-483-5p | 170.7346 | -0.48774 | 0.138011 | -3.5341 | 0.000409 | 0.007716 |
| hsa-miR-320a-3p | 6053.099 | -0.48652 | 0.104609 | -4.6508 | 3.31E-06 | 0.000291 |

| | | | | | | |
|---|---|---|---|---|---|---|
| hsa-miR-331-5p | 45.10024 | -0.48617 | 0.236588 | -2.05493 | 0.039886 | 0.176973 |
| hsa-miR-769-3p | 98.42824 | -0.48285 | 0.160192 | -3.01417 | 0.002577 | 0.027511 |
| hsa-miR-365a-3p | 10095.57 | -0.47934 | 0.117057 | -4.09494 | 4.22E-05 | 0.001393 |
| hsa-miR-365b-3p | 10095.57 | -0.47934 | 0.117057 | -4.09494 | 4.22E-05 | 0.001393 |
| hsa-miR-4521 | 452.6573 | -0.45861 | 0.186377 | -2.46065 | 0.013868 | 0.084167 |
| hsa-miR-212-3p | 92.04929 | -0.44615 | 0.200669 | -2.2233 | 0.026196 | 0.128068 |
| hsa-miR-193b-5p | 247.4601 | -0.4407 | 0.11355 | -3.88109 | 0.000104 | 0.002615 |
| hsa-miR-193b-3p | 442.8027 | -0.43027 | 0.131128 | -3.2813 | 0.001033 | 0.014745 |
| hsa-miR-1843 | 49.54986 | -0.42864 | 0.218517 | -1.96159 | 0.04981 | 0.201144 |
| hsa-miR-423-3p | 4044.027 | -0.40893 | 0.103349 | -3.95679 | 7.60E-05 | 0.002228 |
| hsa-miR-491-5p | 69.89142 | -0.40186 | 0.196505 | -2.04502 | 0.040852 | 0.179245 |
| hsa-miR-106b-3p | 1419.805 | -0.3976 | 0.152669 | -2.60435 | 0.009205 | 0.064803 |
| hsa-miR-193a-5p | 676.5519 | -0.39395 | 0.120339 | -3.27365 | 0.001062 | 0.014752 |
| hsa-miR-138-5p | 3071.165 | -0.39104 | 0.125663 | -3.11182 | 0.001859 | 0.023521 |
| hsa-miR-423-5p | 5085.48 | -0.3881 | 0.12823 | -3.02662 | 0.002473 | 0.027511 |
| hsa-miR-1307-3p | 1490.083 | -0.37841 | 0.145055 | -2.60871 | 0.009088 | 0.064803 |
| hsa-let-7b-3p | 241.2312 | -0.35599 | 0.135292 | -2.63127 | 0.008507 | 0.063261 |
| hsa-miR-7977 | 316.988 | -0.33759 | 0.112126 | -3.01085 | 0.002605 | 0.027511 |
| hsa-miR-29b-1-5p | 130.8471 | -0.32676 | 0.148609 | -2.19876 | 0.027895 | 0.132691 |
| hsa-miR-106a-5p | 1192.932 | -0.27994 | 0.099628 | -2.80986 | 0.004956 | 0.043616 |
| hsa-miR-93-3p | 370.252 | -0.26268 | 0.131403 | -1.99902 | 0.045606 | 0.189131 |
| hsa-miR-362-5p | 306.0997 | -0.25046 | 0.119372 | -2.09811 | 0.035895 | 0.160615 |
| hsa-miR-324-3p | 458.4717 | -0.25018 | 0.099239 | -2.52095 | 0.011704 | 0.075074 |
| hsa-miR-1260a | 1669.818 | -0.23928 | 0.093555 | -2.5576 | 0.01054 | 0.069835 |
| hsa-miR-1260b | 1852.233 | -0.23739 | 0.090677 | -2.618 | 0.008845 | 0.063972 |
| hsa-miR-425-5p | 3517.36 | -0.23441 | 0.109685 | -2.13713 | 0.032588 | 0.147062 |
| hsa-miR-205-5p | 805988.7 | -0.22312 | 0.091036 | -2.45088 | 0.014251 | 0.085504 |
| hsa-miR-22-3p | 10014.19 | -0.2085 | 0.089198 | -2.33751 | 0.019413 | 0.103536 |
| hsa-miR-107 | 3447.842 | -0.20123 | 0.092514 | -2.17513 | 0.02962 | 0.138403 |
| hsa-miR-222-3p | 19687.52 | -0.19445 | 0.081969 | -2.37219 | 0.017683 | 0.097257 |

| | | | | | | |
|---|---|---|---|---|---|---|
| hsa-miR-532-3p | 459.0475 | -0.19271 | 0.095498 | -2.018 | 0.043591 | 0.185614 |
| hsa-miR-20a-5p | 45333.69 | -0.18347 | 0.074189 | -2.47296 | 0.0134 | 0.082269 |
| hsa-miR-23a-3p | 236714.7 | -0.17625 | 0.079229 | -2.22457 | 0.02611 | 0.128068 |
| hsa-miR-128-3p | 5588.19 | 0.164815 | 0.081661 | 2.018286 | 0.043561 | 0.185614 |
| hsa-miR-96-5p | 3104.385 | 0.204514 | 0.074029 | 2.762604 | 0.005734 | 0.048474 |
| hsa-miR-181a-3p | 393.5077 | 0.222626 | 0.11357 | 1.960262 | 0.049965 | 0.201144 |
| hsa-miR-126-3p | 7577.156 | 0.237911 | 0.089854 | 2.64775 | 0.008103 | 0.062005 |
| hsa-miR-130a-3p | 12990.32 | 0.239875 | 0.100997 | 2.375073 | 0.017545 | 0.097257 |
| hsa-miR-205-3p | 1233.262 | 0.257721 | 0.128548 | 2.004866 | 0.044977 | 0.189131 |
| hsa-miR-210-3p | 3664.65 | 0.258687 | 0.119795 | 2.159408 | 0.030819 | 0.142305 |
| hsa-miR-31-3p | 996.2502 | 0.258878 | 0.114577 | 2.259429 | 0.023857 | 0.121119 |
| hsa-miR-190a-5p | 1112.927 | 0.259989 | 0.091336 | 2.846495 | 0.00442 | 0.039878 |
| hsa-miR-30a-3p | 866.2233 | 0.260147 | 0.114152 | 2.278944 | 0.02267 | 0.116213 |
| hsa-miR-148b-3p | 8951.406 | 0.273356 | 0.080741 | 3.385584 | 0.00071 | 0.01103 |
| hsa-let-7f-5p | 248425.6 | 0.280359 | 0.110908 | 2.527844 | 0.011477 | 0.07481 |
| hsa-miR-30e-3p | 1625.034 | 0.28248 | 0.093458 | 3.022546 | 0.002507 | 0.027511 |
| hsa-miR-16-5p | 512593.3 | 0.28267 | 0.117983 | 2.395863 | 0.016581 | 0.095162 |
| hsa-miR-192-5p | 1235.739 | 0.284418 | 0.127386 | 2.232719 | 0.025567 | 0.127355 |
| hsa-miR-452-5p | 1138.978 | 0.295659 | 0.082453 | 3.585774 | 0.000336 | 0.006825 |
| hsa-miR-196b-5p | 6946.906 | 0.306846 | 0.153671 | 1.996772 | 0.04585 | 0.189131 |
| hsa-miR-455-5p | 2449.419 | 0.313645 | 0.094945 | 3.303435 | 0.000955 | 0.014008 |
| hsa-miR-135b-5p | 4506.37 | 0.314087 | 0.116914 | 2.686489 | 0.007221 | 0.056067 |
| hsa-miR-195-5p | 205.9637 | 0.315326 | 0.125581 | 2.510931 | 0.012041 | 0.075661 |
| hsa-miR-30c-1-3p | 129.6073 | 0.323367 | 0.144784 | 2.233446 | 0.02552 | 0.127355 |
| hsa-miR-196a-5p | 7812.09 | 0.323744 | 0.148765 | 2.176217 | 0.029539 | 0.138403 |
| hsa-miR-101-3p | 7502.901 | 0.324681 | 0.117909 | 2.753661 | 0.005893 | 0.048474 |
| hsa-miR-30b-5p | 2523.031 | 0.325547 | 0.10958 | 2.970851 | 0.00297 | 0.030179 |
| hsa-miR-32-5p | 203.2311 | 0.345831 | 0.169297 | 2.042751 | 0.041077 | 0.179245 |
| hsa-miR-7-5p | 12064.13 | 0.364822 | 0.142526 | 2.559696 | 0.010476 | 0.069835 |
| hsa-miR-15b-3p | 1924.135 | 0.36817 | 0.105961 | 3.474592 | 0.000512 | 0.008714 |

| | | | | | | |
|---|---|---|---|---|---|---|
| hsa-miR-203b-3p | 100.8037 | 0.38663 | 0.175478 | 2.203297 | 0.027574 | 0.132691 |
| hsa-miR-1248 | 2562.419 | 0.387728 | 0.176079 | 2.202013 | 0.027664 | 0.132691 |
| hsa-miR-152-3p | 11959.62 | 0.387859 | 0.099741 | 3.888663 | 0.000101 | 0.002615 |
| hsa-miR-218-5p | 1042.18 | 0.388393 | 0.113081 | 3.434644 | 0.000593 | 0.00979 |
| hsa-miR-148a-5p | 101.2574 | 0.390544 | 0.181503 | 2.151726 | 0.031419 | 0.14301 |
| hsa-miR-301a-3p | 1046.184 | 0.401548 | 0.10675 | 3.761561 | 0.000169 | 0.003715 |
| hsa-miR-374a-5p | 3463.037 | 0.408304 | 0.103989 | 3.926409 | 8.62E-05 | 0.002396 |
| hsa-miR-18a-5p | 1338.176 | 0.4165 | 0.162935 | 2.556238 | 0.010581 | 0.069835 |
| hsa-miR-27a-3p | 144185 | 0.420131 | 0.096231 | 4.365841 | 1.27E-05 | 0.000743 |
| hsa-miR-944 | 243.169 | 0.423805 | 0.144104 | 2.940955 | 0.003272 | 0.031411 |
| hsa-miR-12135 | 1378.269 | 0.433031 | 0.18229 | 2.375511 | 0.017525 | 0.097257 |
| hsa-miR-4662a-5p | 102.6433 | 0.442944 | 0.186699 | 2.372501 | 0.017668 | 0.097257 |
| hsa-miR-374a-3p | 186.0495 | 0.444314 | 0.16918 | 2.626274 | 0.008633 | 0.063305 |
| hsa-miR-651-5p | 238.0902 | 0.453897 | 0.177138 | 2.562397 | 0.010395 | 0.069835 |
| hsa-miR-29c-3p | 3704.531 | 0.454434 | 0.112687 | 4.032719 | 5.51E-05 | 0.001712 |
| hsa-miR-27b-3p | 68708.05 | 0.457412 | 0.128807 | 3.551134 | 0.000384 | 0.007501 |
| hsa-miR-6510-3p | 110.8953 | 0.459121 | 0.17739 | 2.588207 | 0.009648 | 0.067026 |
| hsa-miR-3065-3p | 62.46873 | 0.461517 | 0.198606 | 2.323787 | 0.020137 | 0.106323 |
| hsa-miR-3613-5p | 71.22788 | 0.472157 | 0.218881 | 2.157143 | 0.030995 | 0.142305 |
| hsa-miR-20a-3p | 157.0335 | 0.474678 | 0.166789 | 2.845979 | 0.004428 | 0.039878 |
| hsa-miR-106b-5p | 457.5023 | 0.485693 | 0.157915 | 3.075662 | 0.0021 | 0.025204 |
| hsa-miR-126-5p | 1138.997 | 0.488912 | 0.127713 | 3.828207 | 0.000129 | 0.002963 |
| hsa-miR-450b-5p | 141.8594 | 0.495836 | 0.167881 | 2.953496 | 0.003142 | 0.031128 |
| hsa-miR-26b-3p | 108.3206 | 0.498048 | 0.182785 | 2.72478 | 0.006434 | 0.051475 |
| hsa-miR-141-5p | 847.6426 | 0.503215 | 0.113079 | 4.450096 | 8.58E-06 | 0.000566 |
| hsa-miR-32-3p | 55.40411 | 0.520409 | 0.220261 | 2.362696 | 0.018143 | 0.098755 |
| hsa-miR-26a-1-3p | 93.01361 | 0.521194 | 0.170317 | 3.06014 | 0.002212 | 0.025958 |
| hsa-miR-3662 | 44.82965 | 0.521553 | 0.225024 | 2.317766 | 0.020462 | 0.10697 |
| hsa-miR-199b-5p | 137.3796 | 0.522447 | 0.145558 | 3.589283 | 0.000332 | 0.006825 |
| hsa-miR-598-3p | 119.3261 | 0.525121 | 0.151061 | 3.47621 | 0.000509 | 0.008714 |

| | | | | | | |
|---|---|---|---|---|---|---|
| hsa-miR-590-3p | 715.6764 | 0.53279 | 0.113691 | 4.686309 | 2.78E-06 | 0.000291 |
| hsa-miR-29b-3p | 15136.64 | 0.548869 | 0.119448 | 4.595044 | 4.33E-06 | 0.000326 |
| hsa-miR-424-5p | 3183.016 | 0.561839 | 0.136669 | 4.110947 | 3.94E-05 | 0.001393 |
| hsa-miR-301b-3p | 78.16766 | 0.58436 | 0.215089 | 2.716832 | 0.006591 | 0.051941 |
| hsa-miR-140-5p | 174.7749 | 0.592379 | 0.176987 | 3.347011 | 0.000817 | 0.012323 |
| hsa-miR-26a-2-3p | 54.76546 | 0.592657 | 0.235366 | 2.518028 | 0.011801 | 0.075074 |
| hsa-miR-148a-3p | 25429.16 | 0.626995 | 0.184001 | 3.407556 | 0.000655 | 0.010488 |
| hsa-miR-1262 | 40.34907 | 0.636721 | 0.263786 | 2.413782 | 0.015788 | 0.092622 |
| hsa-let-7f-2-3p | 32.14794 | 0.648522 | 0.269324 | 2.407964 | 0.016042 | 0.093077 |
| hsa-miR-3157-5p | 30.58669 | 0.652314 | 0.26021 | 2.506876 | 0.01218 | 0.075661 |
| hsa-miR-301a-5p | 70.90152 | 0.653151 | 0.227237 | 2.874322 | 0.004049 | 0.038176 |
| hsa-miR-30d-3p | 94.78351 | 0.677145 | 0.16233 | 4.171405 | 3.03E-05 | 0.001332 |
| hsa-miR-542-3p | 534.0272 | 0.70017 | 0.131564 | 5.321918 | 1.03E-07 | 1.81E-05 |
| hsa-miR-33a-5p | 124.9803 | 0.704045 | 0.231065 | 3.04696 | 0.002312 | 0.026534 |
| hsa-miR-100-3p | 126.0699 | 0.724725 | 0.245716 | 2.94944 | 0.003184 | 0.031128 |
| hsa-miR-582-5p | 19.30318 | 0.811559 | 0.4 | 2.028901 | 0.042468 | 0.183798 |
| hsa-miR-582-3p | 43.90986 | 0.865093 | 0.224615 | 3.851442 | 0.000117 | 0.002818 |
| hsa-miR-182-3p | 13.39936 | 1.215706 | 0.409246 | 2.970598 | 0.002972 | 0.030179 |
| hsa-miR-1255b-5p | 11.90627 | 1.25666 | 0.456309 | 2.753968 | 0.005888 | 0.048474 |

**Supplementary table 4:** List of unique and common differentially regulated miRNAs by E2 and P4 in VK2 cells (unadjusted $p < 0.05$).

| **13 unique miRNAs in "E2":** | **Log2 FC** | **102 unique miRNAs in "P4":** | **Log2 FC** | **29 common miRNAs in "E2" and "P4":** | **Log2 FC in E2** | **Log2 FC in P4** |
|---|---|---|---|---|---|---|
| hsa-miR-181d-3p | -0.928434279 | hsa-miR-10401-3p | -1.160460613 | hsa-miR-194-3p | -0.812450402 | -0.798340221 |

| | | | | | | |
|---|---|---|---|---|---|---|
| hsa-miR-4804-5p | -0.661601656 | hsa-miR-1246 | -1.063129344 | hsa-miR-1290 | -0.617724484 | -0.885349146 |
| hsa-miR-99a-5p | -0.265811453 | hsa-miR-6883-3p | -1.043694616 | hsa-miR-4485-3p | -0.489904807 | -0.954721424 |
| hsa-miR-9-3p | 0.193553486 | hsa-miR-5701 | -0.824021665 | hsa-miR-12136 | -0.48872638 | -1.157582641 |
| hsa-miR-200c-5p | 0.220748673 | hsa-miR-24-2-5p | -0.784936854 | hsa-miR-181c-5p | -0.479797833 | -0.50697342 |
| hsa-miR-187-3p | 0.464618464 | hsa-miR-3679-5p | -0.643195189 | hsa-miR-7974 | -0.442684128 | -0.607644785 |
| hsa-let-7i-3p | 0.529481783 | hsa-miR-2110 | -0.623220584 | hsa-miR-132-5p | -0.417369882 | -0.522036995 |
| hsa-miR-151b | 0.601185141 | hsa-miR-501-3p | -0.599268039 | hsa-miR-769-3p | -0.416421539 | -0.482845331 |
| hsa-miR-548c-5p | 0.657463547 | hsa-miR-486-5p | -0.56829998 | hsa-miR-483-5p | -0.408983225 | -0.487743449 |
| hsa-miR-548o-5p | 0.657463547 | hsa-miR-132-3p | -0.541933322 | hsa-miR-375-3p | -0.358546043 | -0.711038154 |
| hsa-miR-548am-5p | 0.74388959 | hsa-miR-874-3p | -0.514728766 | hsa-miR-23a-5p | -0.356093208 | -0.552868656 |
| hsa-miR-101-2-5p | 1.001796923 | hsa-miR-331-5p | -0.486170846 | hsa-miR-138-5p | -0.324320649 | -0.391039369 |
| hsa-miR-548au-5p | 1.18634842 | hsa-miR-365a-3p | -0.47934351 | hsa-miR-193b-3p | -0.259935436 | -0.430269401 |
| | | hsa-miR-365b-3p | -0.47934351 | hsa-miR-320a-3p | -0.250374454 | -0.486517161 |
| | | hsa-miR-4521 | -0.45861022 | hsa-miR-423-3p | -0.237484358 | -0.408930436 |
| | | hsa-miR-212-3p | -0.446147986 | hsa-miR-452-5p | 0.195988934 | 0.295659101 |

| | | | | | | |
|---|---|---|---|---|---|---|
| | | hsa-miR-193b-5p | -0.440698313 | hsa-miR-455-5p | 0.210679717 | 0.313645125 |
| | | hsa-miR-1843 | -0.42864097 | hsa-miR-210-3p | 0.261435541 | 0.258687256 |
| | | hsa-miR-491-5p | -0.401857602 | hsa-miR-590-3p | 0.26769946 | 0.532790211 |
| | | hsa-miR-106b-3p | -0.397603069 | hsa-miR-424-5p | 0.281440996 | 0.56183946 |
| | | hsa-miR-193a-5p | -0.393947036 | hsa-miR-205-3p | 0.282419142 | 0.257721072 |
| | | hsa-miR-423-5p | -0.388102675 | hsa-miR-141-5p | 0.320963913 | 0.503214629 |
| | | hsa-miR-1307-3p | -0.378407162 | hsa-miR-20a-3p | 0.34352044 | 0.474677686 |
| | | hsa-let-7b-3p | -0.355988376 | hsa-miR-542-3p | 0.345195355 | 0.700170486 |
| | | hsa-miR-7977 | -0.337594039 | hsa-miR-30d-3p | 0.359012983 | 0.677145329 |
| | | hsa-miR-29b-1-5p | -0.326755994 | hsa-miR-148a-5p | 0.411542804 | 0.390543853 |
| | | hsa-miR-106a-5p | -0.279940053 | hsa-miR-6510-3p | 0.500403664 | 0.459121321 |
| | | hsa-miR-93-3p | -0.262676973 | hsa-miR-3157-5p | 0.646263287 | 0.652314377 |
| | | hsa-miR-362-5p | -0.250456698 | hsa-miR-1255b-5p | 1.0243144 | 1.256660478 |
| | | hsa-miR-324-3p | -0.250177813 | | | |
| | | hsa-miR-1260a | -0.239275588 | | | |

| | | | |
|---|---|---|---|
| | | hsa-miR-1260b | -0.237393694 |
| | | hsa-miR-425-5p | -0.234411259 |
| | | hsa-miR-205-5p | -0.22311767 |
| | | hsa-miR-22-3p | -0.208500642 |
| | | hsa-miR-107 | -0.201230598 |
| | | hsa-miR-222-3p | -0.194445837 |
| | | hsa-miR-532-3p | -0.19271481 |
| | | hsa-miR-20a-5p | -0.183465711 |
| | | hsa-miR-23a-3p | -0.176251477 |
| | | hsa-miR-128-3p | 0.164815149 |
| | | hsa-miR-96-5p | 0.204513946 |
| | | hsa-miR-181a-3p | 0.222626487 |
| | | hsa-miR-126-3p | 0.237910686 |
| | | hsa-miR-130a-3p | 0.239874865 |
| | | hsa-miR-31-3p | 0.258877762 |

| | | | |
|---|---|---|---|
| | | hsa-miR-190a-5p | 0.259988822 |
| | | hsa-miR-30a-3p | 0.260146536 |
| | | hsa-miR-148b-3p | 0.273356012 |
| | | hsa-let-7f-5p | 0.280358602 |
| | | hsa-miR-30e-3p | 0.28247956 |
| | | hsa-miR-16-5p | 0.28267006 |
| | | hsa-miR-192-5p | 0.28441795 |
| | | hsa-miR-196b-5p | 0.306846454 |
| | | hsa-miR-135b-5p | 0.314087346 |
| | | hsa-miR-195-5p | 0.315326268 |
| | | hsa-miR-30c-1-3p | 0.32336656 |
| | | hsa-miR-196a-5p | 0.323744425 |
| | | hsa-miR-101-3p | 0.324680537 |
| | | hsa-miR-30b-5p | 0.325547278 |
| | | hsa-miR-32-5p | 0.345831426 |

| | | | |
|---|---|---|---|
| | | hsa-miR-7-5p | 0.364822179 |
| | | hsa-miR-15b-3p | 0.368170052 |
| | | hsa-miR-203b-3p | 0.386629768 |
| | | hsa-miR-1248 | 0.38772752 |
| | | hsa-miR-152-3p | 0.387859457 |
| | | hsa-miR-218-5p | 0.388392566 |
| | | hsa-miR-301a-3p | 0.401547967 |
| | | hsa-miR-374a-5p | 0.40830422 |
| | | hsa-miR-18a-5p | 0.416500079 |
| | | hsa-miR-27a-3p | 0.420131053 |
| | | hsa-miR-944 | 0.42380457 |
| | | hsa-miR-12135 | 0.433030707 |
| | | hsa-miR-4662a-5p | 0.442944244 |
| | | hsa-miR-374a-3p | 0.444314067 |
| | | hsa-miR-651-5p | 0.453896874 |

| | | | |
|---|---|---|---|
| | | hsa-miR-29c-3p | 0.454434145 |
| | | hsa-miR-27b-3p | 0.457412403 |
| | | hsa-miR-3065-3p | 0.461517119 |
| | | hsa-miR-3613-5p | 0.472156762 |
| | | hsa-miR-106b-5p | 0.48569328 |
| | | hsa-miR-126-5p | 0.488911946 |
| | | hsa-miR-450b-5p | 0.49583615 |
| | | hsa-miR-26b-3p | 0.498048441 |
| | | hsa-miR-32-3p | 0.520408519 |
| | | hsa-miR-26a-1-3p | 0.521193789 |
| | | hsa-miR-3662 | 0.521552914 |
| | | hsa-miR-199b-5p | 0.522447414 |
| | | hsa-miR-598-3p | 0.525120517 |
| | | hsa-miR-29b-3p | 0.548869113 |
| | | hsa-miR-301b-3p | 0.584359674 |

| | | | |
|---|---|---|---|
| | | hsa-miR-140-5p | 0.592378952 |
| | | hsa-miR-26a-2-3p | 0.592657043 |
| | | hsa-miR-148a-3p | 0.626995056 |
| | | hsa-miR-1262 | 0.636721401 |
| | | hsa-let-7f-2-3p | 0.648521829 |
| | | hsa-miR-301a-5p | 0.653151154 |
| | | hsa-miR-33a-5p | 0.704044888 |
| | | hsa-miR-100-3p | 0.724724666 |
| | | hsa-miR-582-5p | 0.811559347 |
| | | hsa-miR-582-3p | 0.865092701 |
| | | hsa-miR-182-3p | 1.215705571 |
